# PERSEUcpp: A machine learning strategy to predict cell-penetrating peptides and their uptake efficiency

**DOI:** 10.1101/2025.04.07.647598

**Authors:** Rayane Monique Bernardes-Loch, Gustavo de Oliveira Almeida, Igor Teixeira Brasiliano, Wagner Meira, Douglas E V Pires, Maria Cristina Baracat-Pereira, Sabrina de Azevedo Silveira

**Affiliations:** Department of Biochemistry and Molecular Biology, Universidade Federal de Viçosa, 36570-900 Viçosa-MG, Brazil; Department of Computer Science, Universidade Federal de Viçosa, 36570-900, Viçosa - MG, Brazil; Department of Computer Science, Universidade Federal de Minas Gerais, Antônio Carlos Av. 6627, 31270-901, Belo Horizonte - MG, Brazil; School of Computing and Information Systems, University of Melbourne, Parkville 3052, Australia

**Keywords:** Cell-penetrating Peptide, Machine Learning, Uptake Efficiency, Interpretable Machine Learning, CPP Prediction

## Abstract

**Motivation:** Cell-penetrating peptides (CPPs) are promising tools for transporting therapeutic molecules into cells without damaging the cellular membrane. These peptides serve as efficient drug delivery systems, capable of carrying diverse biologically active substances while exhibiting low cytotoxicity compared to non-native molecules. However, identifying CPPs through experimental methods is expensive and time-consuming, making computational strategies an attractive alternative due to their cost-effectiveness and scalability.

**Results:** This study introduces PERSEUcpp, a machine learning strategy designed to identify CPPs. Based on descriptors including physicochemical and structural properties as well as atomic composition, our strategy employs the Extremely Randomized Trees to predict CPPs and their uptake efficiency. The first stage was developed using a balanced dataset of 967 CPPs and non-CPPs, applying a 10-fold cross-validation scheme. Two independent datasets were utilized for validation. The CPP predictor achieved superior results compared to state-of-the-art methods, with MCC 0.854, Recall 0.860, and AUC 0.984. The second stage, focused on efficiency prediction, was trained on a balanced dataset of 140 CPPs and non-CPPs, also using a 10-fold cross-validation scheme, and validated with an independent dataset. The efficiency predictor achieved competitive results, with Recall 0.761 and AUC 0.690. PERSEUcpp is interpretable, offering insights into the key features enabling peptides to penetrate cells effectively. We anticipate that PERSEUcpp will be a valuable tool for advancing the design and application of CPPs in drug delivery and biomedical research.

**Availability and Implementation:** All the source code and datasets are available at GitHub: https://github.com/goalmeida05/PERSEU.

## Introduction

One of the most remarkable achievements of medicine in the twentieth century was the discovery of antibiotics [1, 2]. Their use in clinical routine for treating bacterial infections significantly reduced patient mortality. During the so-called “Golden Age of antibiotics”, there were significant discoveries of new antibacterial drugs [2].

However, despite the existence of these drugs, bacterial resistance began to be observed. The Center for Disease Control (CDC) estimates that nearly 2 million deaths occur in the United States due to bacterial resistance to first-line antibiotics [3]. Almost all known bacterial species have some gene that leads to antibiotic resistance [4]. Bacterial species resistant to antibiotics were also detected in several environmental sites, such as surface water bodies, agricultural soils, and animal manure [5]. Given this scenario, it is imperative to search for new molecules with potential to circumvent antibiotic resistance.

Cell-penetrating peptides (CPPs) are strong contenders in addressing this problem. CPPs are characterized by a short chain of amino acids (usually up to 30 residues in length), partially hydrophobic, usually with a net positive charge, as they are rich in arginine and lysine and show a high isoelectric point [6]. CPPs are known for their ability to transport various therapeutic molecules into cells, including antibiotics [7– 9]. This can potentially enhance the treatment efficacy and overcome some resistance mechanisms [6]. An example is the peptide trans-activator of transcription from HIV (TAT), which is well-known for its ability to transport therapeutic molecules into cells [10]. Additionally, using CPPs to transport antibiotics directly into resistant bacteria can reduce the necessary antibiotic concentration, minimizing side effects and decreasing the development of additional resistance [6].

Determining a peptide’s ability to penetrate cells through *in vitro* and *in vivo* experimental methods is expensive and time-consuming. Therefore, computational methods such as machine learning (ML) [11–13], molecular modeling tools [14– 17] and probabilistic models [18, 19] have been applied to identify the most crucial characteristics for analysis. The goal is to facilitate and enhance the accuracy of predictions, allowing for a more streamlined approach without the need for extensive experimental assays.

In an effort to improve the classification of CPPs, recent studies [20–25] have investigated the integration of various descriptors from sequence, structure, and physicochemical properties. In Supplementary Section 1 we briefly review some representative examples of the mentioned studies.

Despite significant progress of predictive methods for cell-penetrating peptides, some limitations persist. An important drawback is the lack of experimentally validated negative data. In general, models utilize artificially generated negative examples based on a small sample of experimentally validated non-CPPs. This approach does not guarantee that a positive example cannot be generated, which may lead to imprecise classifications. High computational cost is also a significant issue, particularly for models like the k-skip-n-gram, which, while effective in capturing spatial information, can become inefficient and resource-intensive when applied to large datasets. Moreover, deep learning approaches require substantial computational resources and large training datasets, which might not be accessible to all researchers. These models also tend to lose interpretability, making it challenging to understand the underlying factors contributing to their predictions.

To overcome some of these challenges we propose PERSEUcpp, an interpretable strategy that leverages features related to structural, physicochemical properties and atomic composition. We propose a novel method that models each experimentally identified CPP as a feature vector containing the above mentioned properties. Next, the resulting matrix is fed into an Extremely Randomized Trees (ERT) algorithm to assess whether a given peptide possesses a cell-penetrating function. A similar process was conducted to train the efficiency classifier. Our results demonstrate the superior predictive capability of PERSEUcpp and highlight the biological significance of the selected descriptors for CPP prediction. All the source code and datasets are freely available at https://github.com/goalmeida05/PERSEU.

## Materials and methods

This section details PERSEUcpp, our supervised learning strategy based on structural, physicochemical and atomic composition descriptors to predict cell-pentrating peptides. Here we explain how data was collected and prepared, the encode of CPPs as feature vectors, the supervised learning step as well as the evaluation strategy. Figure 1 presents a workflow that summarizes the main steps of our method.

**Fig. 1.**
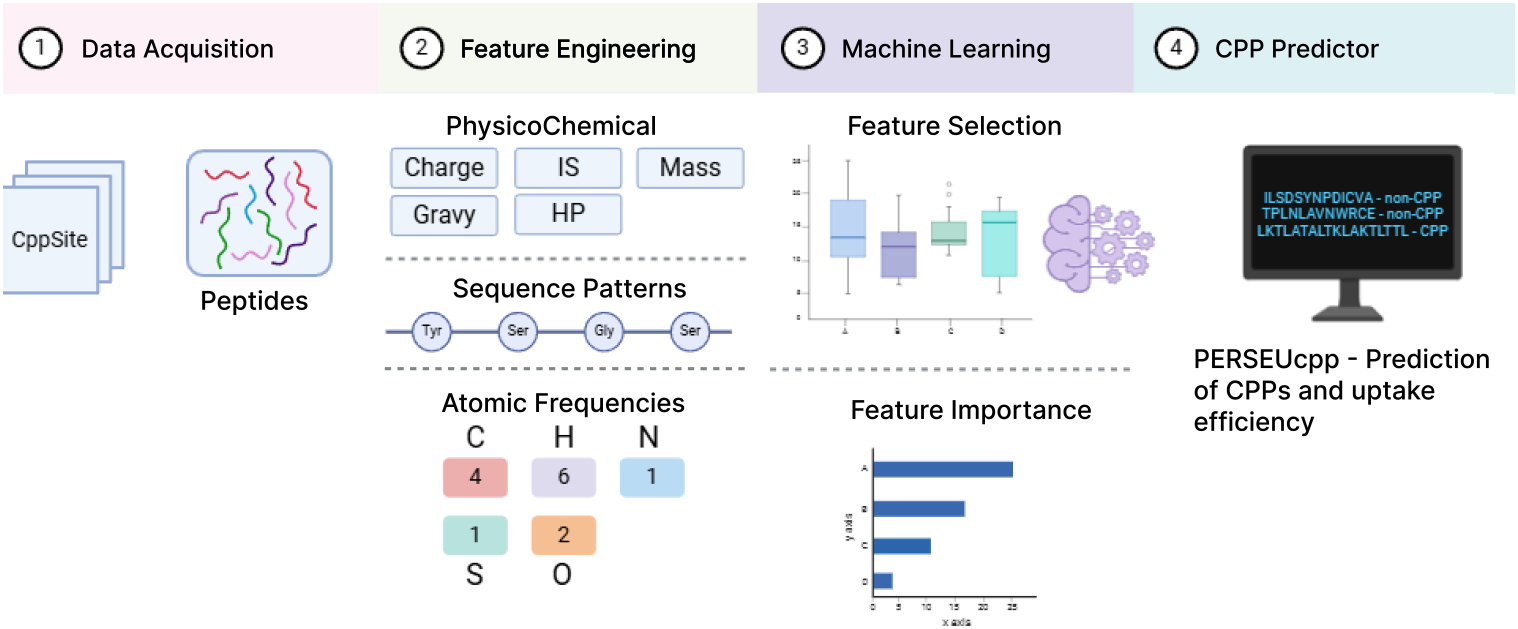
PERSEUcpp workflow. The workflow is segmented into four blocks. **1** Data Acquisition, peptides sequences was acquired from CppSite2.0 and MLCPP2.0; **2** Feature engineering, which involved calculating three classes of features: (i) Physicochemical properties, (ii) Sequence Patterns, which calculate frequencies of residues in the peptide’s chain; (iii) Atomic Frequencies, which calculate frequencies of each atom in peptide’s chain; **3** were then utilized for training and testing models via supervised learning, with feature selection conducted for model optimization. **4** best-performing model was implemented, and all the source code is freely available at GitHub.

### Dataset preparation

Two datasets were used to train our method, one for CPP classification and the other for CPP uptake efficiency classification. Three independent datasets were used in the evaluation to demonstrate its generalization capabilities and to compare PERSEUcpp with alternative methods. In the literature, there are datasets that are a composition of existing ones. Therefore, below, we detail all the datasets that were used in this work, directly or indirectly.

1. **CPPSite2.0:** Composed of 1,883 unique CPPs. CPPsite2.0 is a continuously updated dataset of experimentally validated CPPs. It was used to augment our training dataset, which we named *PerseuCPP training dataset*.
2. **MLCPP2.0 training dataset:** Originally composed of 573 experimentally validated CPPs from CPPSite2.0 and 573 non-CPPs. The non-CPPs share less than 15% sequence identity with non-CPPs and with CPPs [24] in this set.
3. **PerseuCPP training dataset:** Our dataset is composed of the MLCPP2.0 training set, augmented with additional positive sequences from CPPSite2.0. To balance the number of newly added positive sequences with negatives, we downloaded data from several previously published methods [20–25], then excluded any sequences that appeared in the test sets (MLCPP2.0 independent dataset, CPP924 independet dataset, CPP1708 test dataset) and removed duplicates. The resulting dataset contained natural peptides with 967 CPPs and 944 non-CPPs, which we used to build our classification model.
4. **MLCPP2.0 efficiency training dataset:** Composed of 140 peptides with low and 140 CPPs with high uptake efficiency. This dataset was used to build our model for CPP uptake efficiency classification task, and we named it *train-dataset-2*.
5. **MLCPP2.0 independent dataset:** Composed of 157 experimentally validated CPPs and 2184 non-CPPs. These data were used to compare PERSEUcpp with competitors.
6. **MLCPP2.0 efficiency independent dataset:** Composed of 40 CPPs with low and 40 CPPs with high uptake efficiency. The purpose of this dataset was to compare PERSEUcpp with competitors [24].
7. **CPP924 independent dataset:** A high-quality dataset consisting of 462 unique natural CPPs and non-CPPs experimentally validated. The sequence similarity between any two sequences in the positive group was less than 20% [23].The purpose of this dataset was to compare PERSEUcpp with competitors.
8. **CPP1708 dataset:** Composed of 854 experimentally validated CPPs and 854 non-CPPs [27]. The dataset was generated by deduplicating sequences from multiple sources using CD-HIT at a 70% similarity threshold and balancing the number of positive and negative samples through random sampling. Stratified sampling based on molecular weight was applied to create an 80/10/10 split for training, validation, and testing.

### Feature engineering

In the first part of this study, a set of 8,831 features was calculated, normalized by the chain length of each peptide sequence. These descriptors included *Amino Acid Composition* (AAC), *Atomic Composition, DiPeptide Composition* (DPC), *TriPeptide Composition (TPC), Composition of k-spaced Amino Acid Group Pairs* (CKSAAGP), *Physicochemical properties* such as Grand Average of Hydropathy (GRAVY) index, Molecular Mass, Isoelectric Point, Hydrophobicity, Net Charge.

We employed the *AAC*, a straightforward yet widely utilized feature descriptor of considerable importance for peptide composition. This descriptor computes the frequency of each of the 20 amino acids occurring in peptide sequences and their respective normalization within each chain.

For *DPC*, all possible combinations of the 20 amino acids, taken pairwise, were computed. Also, we used DPC normalized by the peptide length. For *TPC*, all possible combinations of the 20 amino acids taken in groups of three were calculated. Additionally, we used TPC normalized by the length of the peptide.

For the *CKSAAGP*, we calculated the frequency of pairs of amino acid groups separated by *K* residues in the peptide sequence. Amino acids were grouped based on their physicochemical properties, such as charge, polarity, and hydrophobic groups. This feature captures proximal interactions between amino acid groups that can significantly influence the peptide structure and function. Additionally, the CKSAAGP descriptors were normalized by the peptide length to ensure consistency across sequences of varying lengths.

The *physicochemical properties*: GRAVY index, Molecular Mass, Isoelectric Point, Hydrophobicity and Net Charge were calculated using Biopython library [28]. Finally, the *Atomic Composition* was calculated by determining the carbon, hydrogen, nitrogen, oxygen, and sulfur quantities and normalized by chain length.

Due to the large number of features, an exploratory data analysis was conducted using the Mann-Whitney statistical test to determine whether each feature showed a significant difference (p-value *<* 0.05) between the CPP and non-CPP groups. As a result, the initial set of 8,831 features was reduced to 1,137. This is available in Supplementary Subsection 2.1.

### Evaluation strategy

#### Model training

We applied Extremely Randomized Tree (ERT), Random Forest (RF), XGBoost (XGB), Support Vector Machine (SVM), and Multilayer Perceptron (MLP) algorithms to our training dataset (*PerseuCPP training dataset:*) to identify the one that would produce the best results. Additionally, a Singular Value Decomposition (SVD) was applied to perform dimensionality reduction and the model performance was evaluated considering up to 100 singular values. We conducted a 10-fold cross-validation yielding an average performance that reflects the model stability across different training and testing data configurations.

#### Comparison with alternative methods

We evaluate PERSEUcpp and compare it with state-of-the-art methods using the same set of metrics as the competitors such as Sensitivity (SN), Specificity (SP), Accuracy (ACC), Matthews correlation coefficient (MCC), F1, and Area Under the Receiver Operating Characteristic Curve (AUC). The presented metrics are explained and mathematically defined in the subsection *Comparison with alternative methods* in Supplementary Subsection 2.2.

## Results and discussion

To demonstrate our method’s ability to predict CPPs, we conducted a comprehensive set of experiments. First, we evaluated five different algorithms (ERT, RF, XGB, SVM and MLP) to determine which would be the best for building the model. After selecting the best algorithm, we compared our model with five other predictors, including the state-of-the-art. Additionally, we devised an efficiency predictor for CPPs and compared it with the state-of-the-art method.

### PERSEUcpp cross-validation results

First, we assessed which algorithm (ERT, RF, XGB, SVM, and MLP) would perform best in the classification of CPPs. We applied the same training matrix for each algorithm and all of them were evaluated using a 10-fold cross-validation. We tested different configurations of the dataset, keeping the same data and features but applying different treatments to each. The results of these analyses are provided in the Supplementary Section 3.1.

The Extremely Randomized Trees algorithm exhibited the best performance and was crucial for selecting the most informative features, thus highlighting the importance of each descriptor. From the original set of 1,137 features, we reduced the dimensionality to 522, as detailed in Table S12 of Supplementary Subsection 3.2.

### PERSEUcpp results compared with state-of-the-art methods

#### CPP924 validation dataset

In this section, the performance of PERSEUcpp was compared with four CPP predictors: CPPred-RF, TargetCPP, StackCPPerd, and SiameseCPP. The comparisons were conducted using the CPP924 validation dataset to evaluate the performance of our method regarding existing predictors. This dataset is balanced between CPPs and non-CPPs, comprising 924 natural peptides, all of which have been experimentally validated.

SiameseCPP is a deep-learning architecture that learns discriminative features directly from peptide sequences [25]. These features were fused with contrastive features to achieve good classification performance. The method was built with CPP924 dataset, in which 80% was used to train and 20% to test and was compared with competitors. Despite generally presenting good results, the model lacks interpretability and practical biological insights.

According to [29], building and testing a classification model with 80% of the data used to train and 20% to test is less robust than cross-validation, and there is a possibility that the results obtained in this way could be overestimated. However, for fair comparison, we used the same partition with 20% of the dataset to test our method against competitors, as it was the same configuration used by the then state-of-the-art method. Table 1 presents the comparison between PERSEUcpp and the competing methods on CPP924. Our strategy presents results that are superior to SiameseCPP on the CPP924 dataset.

**Table 1.** Comparison of PERSEUcpp results with competing methods on dataset CPP924.

| Method | MCC | ACC | SN | SP | AUC |
| --- | --- | --- | --- | --- | --- |
| CPPred-RF | 0.831 | 0.916 | 0.905 | 0.926 | - |
| TargetCPP | 0.871 | 0.935 | 0.943 | 0.936 | - |
| StackCPPred | 0.890 | 0.945 | 0.942 | 0.948 | - |
| SiameseCPP | 0.923 | 0.961 | 0.959 | 0.964 | - |
| PractiCPP | 0.913 | 0.956 | 0.943 | 0.971 | - |
| **PERSEUcpp** | **0.940** | **0.970** | **0.961** | **0.978** | 0.975 |

The bold values denote the best performance among state-of-the-art methods. The AUC values were not informed by the authors.

PractiCPP [35] employs hard negative sampling to enhance decision boundaries and utilizes three feature types: sequential features from amino acid sequences, local features derived from Morgan fingerprints (which capture the chemical environment around each atom [36]), and pretrained embeddings from the ESM-2 language model. PractiCPP outperforms state-of-the-art models like SiameseCPP and MLCPP2.0 on both balanced and imbalanced datasets. On the other hand, our model outperforms PractiCPP on the CPP924 dataset, Table 1. Nevertheless, it was not possible to undertake a fair comparison using the imbalanced dataset, as it was not provided alongside the source code in the PractiCPP article.

#### MLCPP2.0 validation dataset

In this section, the performance of PERSEUcpp was compared with four CPP predictors: C2Pred, BchemRF, MLCPP2.0, and SiameseCPP. For this purpose, the MLCPP2.0 validation dataset was employed, which was likewise utilised as a benchmark by the aforementioned four methods. This dataset comprises 157 CPPs (experimentally validated) and 2,184 non-CPPs (generated automatically).

The results on Table 2 demonstrate that PERSEUcpp outperforms the methods C2Pred, BchemRF, and MLCPP 2.0, as well as the SiameseCPP. This superiority highlights the effectiveness of PERSEUcpp and its potential practical application in CPP predictions.

**Table 2.**
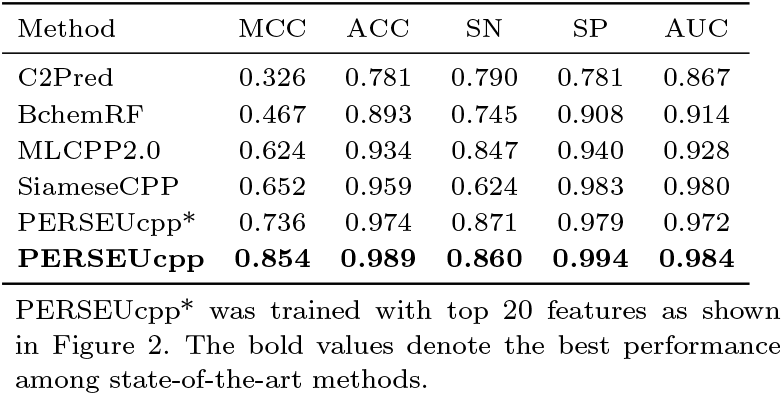
Comparison of PERSEUcpp results with competing methods on dataset MLCPP2.0.

#### CPP1708 validation dataset

In this section, the performance of PERSEUcpp was compared with three CPP prediction methods: CellPPD, C2Pred, MLCPP2.0, and GraphCPP. This dataset comprises a diverse set of 90 CPPs and 98 non-CPPs, designed to test the generalization ability of predictive models.

GraphCPP [27] leverages graph neural networks (GNNs) to predict cell-penetrating peptides (CPPs) by modeling peptide sequences as molecular graphs. This method captures intricate relationships between amino acids and their physicochemical properties. By integrating node and edge features, GraphCPP provides robust embeddings that enhance the accuracy of CPP prediction.

To evaluate the robustness of our model, we also trained PerseuCPP on the CPP1708 training set, which contains 854 experimentally validated CPPs and 854 non-CPPs; the results are shown in Table 3 under the label “PerseuCPP*” Additionally, to capture higher-level semantic information for both our main training dataset (“PerseuCPP**”) and the CPP1708 training set (“PerseuCPP***”), we generated protein language embeddings using the ESM1-T6-43M-UR50S model by averaging the token representations from its sixth layer, thus producing a fixed-length vector for each sequence. While this approach considerably increased the dimensionality of the feature space (from 522 to 1289 features, including 767 ESM-derived descriptors), we applied Singular Value Decomposition (SVD) to iteratively test and determine the optimal number of components. Ultimately, a reduction to 43 components provided the best predictive performance on the test set, as reported in Table 3.

**Table 3.**
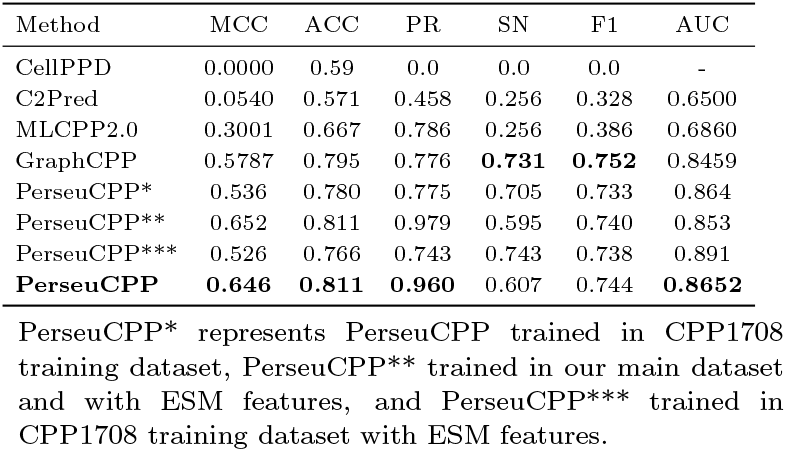
Comparison of PERSEUcpp results with competing methods on dataset CPP1708 Independent dataset.

The results presented in Table 3 demonstrate that PERSEUcpp showing compatible or superior results in comparison with the state-of-the-art competitor, as presented in Table 3, showcasing its robust prediction capability even under challenging conditions.

### Feature importance

The first part of this study was focused on the effects of combining physicochemical, structural, and atomic characteristics on the classification of CPPs. The algorithm Extra Trees was instrumental in this, providing insights into the importance of each feature. Figure 2 presents the top 20 descriptors, underscoring their relevant role in the classification process.

**Fig. 2.**
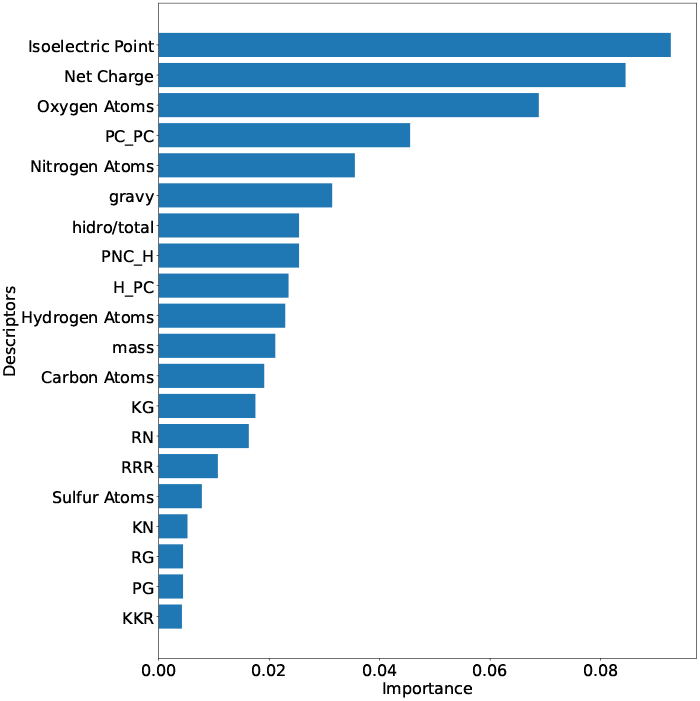
Top 20 independent descriptors for CPP classification identified by Extremely Randomized Trees algorithm. The abbreviations PC, NPC, and H represent the amino acid groups: charged, polar non-charged, and hydrophobic, respectively.

The *Isoelectric Point* and *Net Charge* are descriptors in the group of physicochemical properties and are highly influential in the characterization of CPPs. In Figure 3, it was confirmed that the analyzed CPPs predominantly presented values of isoelectric point greater than seven (*pI >* 7), thus being cationic in their vast majority under physiological conditions. According to the literature, the cell membrane is primarily composed of phospholipids, that could shown phosphate groups and substituent groups negatively charged in their polar heads [30], facilitating the interaction of CPPs with cell membranes, especially in bacteria, confirming the relevance of these characteristics in the classification of CPPs.

**Fig. 3.**
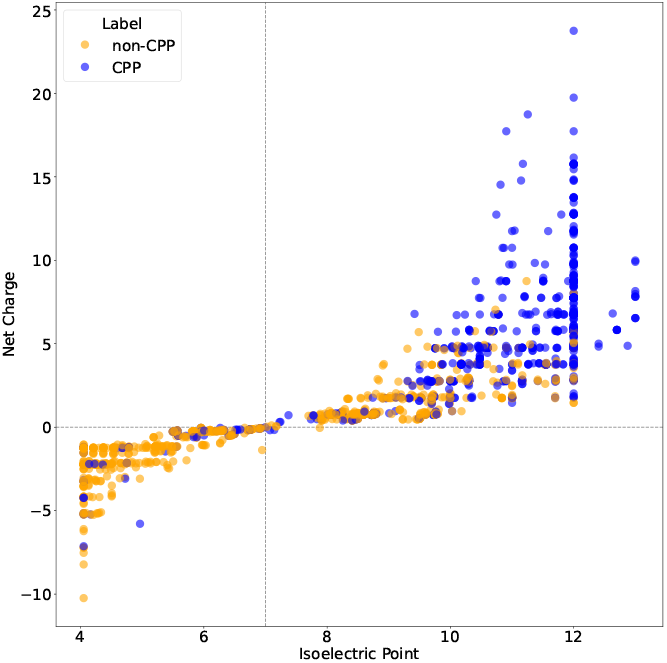
Scatter plot of CPPs and non-CPPs based on isoelectric point and net charge on training dataset. CPPs are predominantly cationic peptides, showing values of isoelectric point superior to 7 under physiological conditions, while a large number of non-CPP are anionic peptides, with values of isoelectric point inferior to 7. The dotted line at *x* = 7 indicates the isoelectric point, highlighting where the charge transition begins.

These top 20 features in Figure 2 are sufficient for the classification of CPPs, even though there is a low presence of non-CPP peptides with high isoelectric points (cationic) and CPPs with isoelectric points below seven and a negative charge, as shown in Figure 3. The model trained exclusively with these 20 features demonstrates reasonable performance compared to competitors as shown in Table 2. However, incorporating new features is necessary to ensure better data separation.

Studies suggest that a balance among physicochemical characteristics is critical for a peptide to interact with cellular membranes [31, 34]. This behavior is observed in the model, as besides *Isoelectric Point* and *Net Charge, Gravy, Hydrophobicity*, and *Mass* occupy positions 6, 7, and 11 in importance in Figure 2.

Arginine (R) and lysine (K) are among the residues relevant for the effectiveness of cell-penetrating peptides (CPPs), since they are cationic. Arginine-rich CPPs hold promise for delivering therapeutic macromolecules such as peptides, proteins, and nucleic acids into cells [32]. The significance of arginine in CPPs is underscored by its involvement in electrostatic interactions and its contribution to the binding affinity of CPPs to cell membranes [32]. Furthermore, lysine-rich peptides have demonstrated higher cell-penetrating activity compared to other peptides [33]. It is, therefore, reasonable that the model identified arginine and lysine as critical features for classification, as shown in Figure 2. These amino acids are highlighted in dipeptides *KG, RN, KN*, and *RG* at positions 13, 14, 17, and 18, respectively, as well as in tripeptides *RRR* and *KKR* at positions 15 and 20. Based on this, we also analyzed the presence of each amino acid in peptide chains. Corroborating this information, Figure 4 shows the frequency of each amino acid in peptide chains, with arginine and lysine appearing in greater abundance in chains experimentally classified as CPPs.

**Fig. 4.**
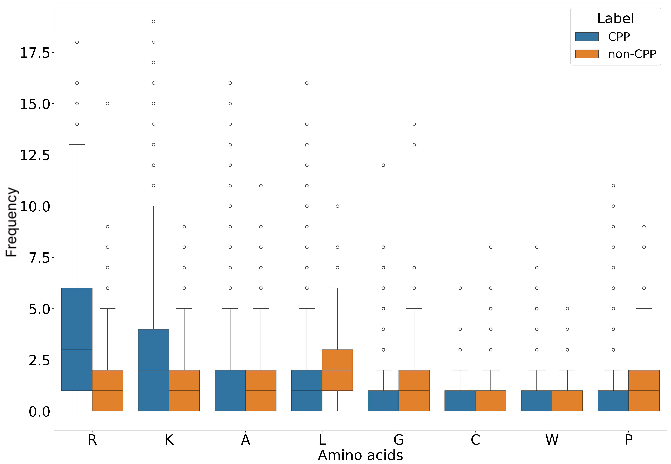
Boxplot comparing the quantity of different amino acids. The x-axis represents the amino acids, while the vertical y-axis indicates the frequency of these amino acids in each sequence. The boxes display the median, quartiles, and outliers for each group.

The utility of dipeptide composition extends to predicting subcellular protein localization [37], identifying bacterial toxin proteins [38], and designing highly efficient cell-penetrating peptides [20]. Particularly in the search for CPPs, dipeptide composition has been employed to differentiate between CPPs and non-CPPs [20]. As can be observed in Figure 5, the CPPs/non-CPPs ratio for the dipeptides *KG, RN, KN, RG*, and *PG* suggests that the presence of *KN* and *PG* is more common in CPPs, while *KG, RN*, and *RG* are more present in non-CPPs.

**Fig. 5.**
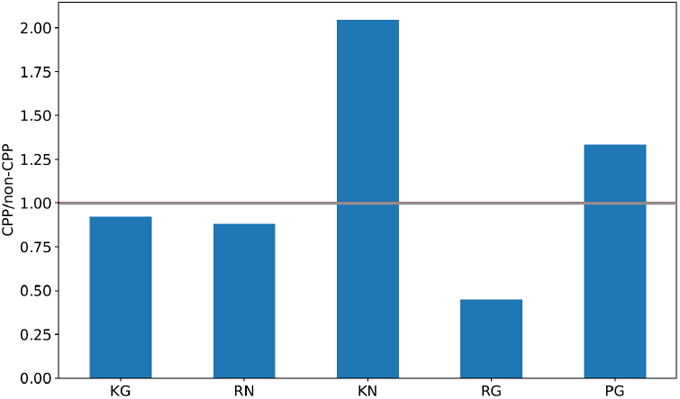
The ratio of the frequencies of the features KG, RN, KN, RG and PG between the groups CPP and non-CPP. Each bar in the chart represents the ratio of the frequencies of a specific feature, calculated as frequency of the feature in the CPP group divided by the frequency of the same feature in the non-CPP group. Values above 1 suggest that the frequency of the feature is higher in the CPP group. Values below 1 suggest that the frequency of the feature is higher in the non-CPP group.

Each amino acid residue comprises a certain amount of carbon (nC), hydrogen (nH), nitrogen (nN), oxygen (nO), and sulfur (nS) in its composition. In this work, the frequency of these atoms normalized by chain length are used as descriptors.It is observed in Figure 2 that the descriptors *Oxygen Atomsand* and *Nitrogen Atoms* play an important role in classification.The relationship between the number of nitrogen and oxygen atoms and the cell-penetrating capabilities of peptides has been investigated, highlighting the significance of these atoms in enhancing peptide function [39, 40]. Specifically, argininerichpeptides, which contain multiple nitrogen atoms in their guanidinium groups, have been shown to penetrate cell membranes effectively. This is primarily due to their ability to form hydrogen bonds and engage in electrostatic interactions with negatively charged cell membrane components, such as phospholipids and glycosaminoglycans [39].

Moreover, the presence of oxygen atoms in specific peptides,often in the form of hydroxyl groups or carbonyl groups, can contribute to their amphipathic nature, which is essential forthe membrane translocation process. These groups facilitate interactions with the membrane’s hydrophilic and hydrophobic regions, thereby improving the peptide ability to cross the lipid bilayer [40].

It is worth noting that the amount of nitrogen is higher in CPPs compared to non-CPPs, whereas the amount of oxygen is greater in non-CPPs as shown in Figure 6. This result is justified by the radicals of amino acid residues that may contain ionizable groups, becoming positive by the protonation of the amino groups (in the presence of nitrogen), or becoming negative by the deprotonation of the carbonyl or hydroxyl groups (in the presence of oxygen) at physiological pH. Thus,cationic peptides are rich in nitrogen, while anionic peptides are rich in oxygen.

**Fig. 6.**
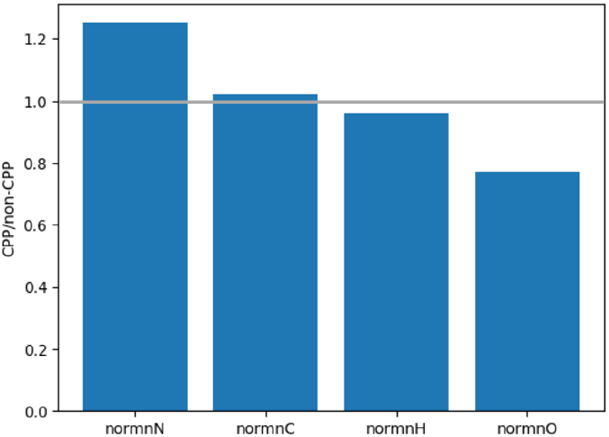
The ratio of the normalized means of the features normnN,normnO, normnH, and normnC between the groups CPP and non-CPP. Each bar in the chart represents the ratio of the normalized mean of a specific feature, calculated as the normalized mean of the feature in the CPP group divided by the normalized mean of the same feature in the non-CPP group. Values above 1 suggest that the normalized mean of the feature is higher in the CPP group. Values below 1 suggest that the normalized mean of the feature is higher in the non-CPP group.

As shown in Figure 2, the features *PC_PC, PNC_H* and *H_PC* as part of the CKSAAGP (Composition of k-spaced Amino Acid Group Pairs) with amino acid distance of S=1also had a strong influence on the classification. This feature captures the frequency of pairs of amino acid groups separated by one residue, reflecting proximal interactions that can directly influence the structure and function of the peptides.Specifically, interactions between positive charges with positive charges (PC_PC), uncharged polar groups with hydrophobic ones (NPC H), and hydrophobic with positive charges (H_PC) are described in the literature as relevant for the cell penetration capability [8]. These interactions affect crucial properties such as cell membrane affinity, conformational stability, and solubility, contributing significantly to the efficiency of peptide penetration.

Figure 7 illustrates the relative contribution of different descriptor groups in the classification of cell-penetrating peptides by PERSEUcpp. The analysis reveals that *dipeptides* and *tripeptides* have significant influence in the classification task. However, when analyzed in isolation, i.e., descriptor by descriptor, they appear to have little influence, as could be expected, considering that there are 20^2^ dipeptides and 20^3^ tripeptides. Other features have significant impact even when considered in isolation. For instance, the *isoelectric point*, which is part of the physicochemical group of descriptors, presents an individual contribution of nearly 10%. Also, the other physicochemical descriptors, such as *net charge*,*hydrophobicity, gravy* and *mass*, also play an important role, especially for cationic peptides that interact with anionic regions of phospholipids on the cell surface [30]. Additionally, *atomic composition* showed a moderate contribution, suggesting that specific elements, like nitrogen and oxygen, influence electrostatic interactions and hydrogen bonding with membrane components [39, 40]. Overall, this analysis of descriptors segmented in categories indicates that each group has a relevant contribution for the prediction of CPPs, emphasizing the need for features that perceive physicochemical properties, composition of residues as well as some context (as captured by di and tripeptides), as all of these are connected with CPPs ability to deliver therapeutic molecules.

**Fig. 7.**
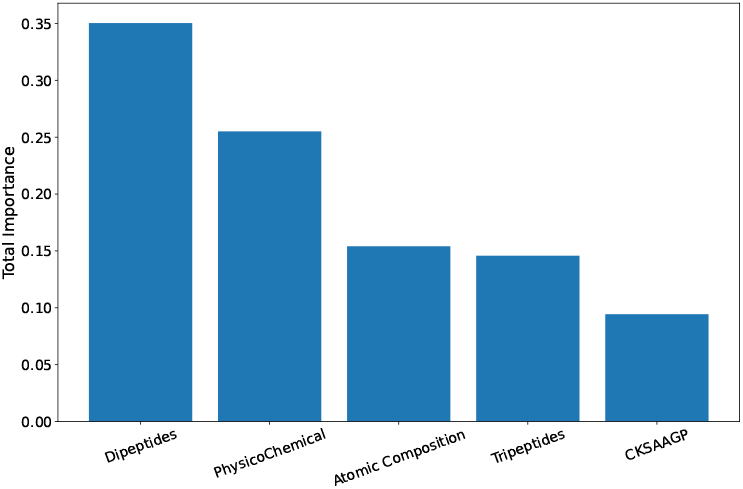
Relative contribution of different groups of descriptors in the classification of CPPs. Each bar represents the contribution of the feature group, in which dipeptides represent groups of two amino acids and tripeptides represent groups of three. The physicochemical group includes features such as GRAVY index (Grand Average of Hydropathicity), Molecular Mass, Isoelectric Point, Hydrophobicity, and Net Charge. Atomic Composition refers to the atoms of nitrogen, carbon, hydrogen, oxygen, and sulfur. CKSAAGP represents the Composition of k-spaced Amino Acid Group Pairs.

In Suplementary Subsection 3.2, Figure S7, we presents the t-distributed Stochastic Neighbor Embedding (t-SNE) visualization, a dimensionality reduction strategy used to visualize complex, high-dimensional data, showed a significant separation between CPPs and non-CPPs.

### Incorrect predictions

Considering the independent datasets CPP924, MLCPP2.0 and CPP1708, our model incorrectly predicts 98 in 2699 sequences, with 5 from CPP924, 35 from MLCPP2.0 and 58 from CPP1708. Table 4 shows the number of incorrect predictions for each dataset considering CPPs and non-CPPs.

**Table 4.** Incorrect predictions in PERSEUcpp. Freq. is the number of sequences in CPP and nonCPP subsets, CP is correct predictions and IP is incorrect predictions.

| Base | Label | Freq. | CP | IP |
| --- | --- | --- | --- | --- |
| CPP924 | CPP | 92 | 88 | 4 |
|  | non-CPP | 93 | 92 | 1 |
| MLCPP | CPP | 157 | 150 | 7 |
|  | non-CPP | 2170 | 2142 | 28 |
| CPP1708 | CPP | 79 | 50 | 29 |
|  | non-CPP | 96 | 67 | 29 |

To illustrate, here we briefly discuss the descriptors *isoelectric_point* and *net_charge*, which are the most important features taken separately. In Figure 8A in Supplementary Section 4, the median *isoelectric_point* of positive train sequences has a value close to the median *isoelectric_point* of the wrongly predicted sequences in Figure 8B. A similar behavior is observed when analyzing the *net_charge*, as in Figure 8C the median *net charge* of positive train sequences has a value close to the median *net charge* of wrongly predicted sequences in Figure 8D. Thus, it seems that what causes confusion in our model is that the negative wrongly predicted sequences (non-CPPs predicted as CPPs), have descriptors similar to the positive train sequences.

**Fig. 8.**
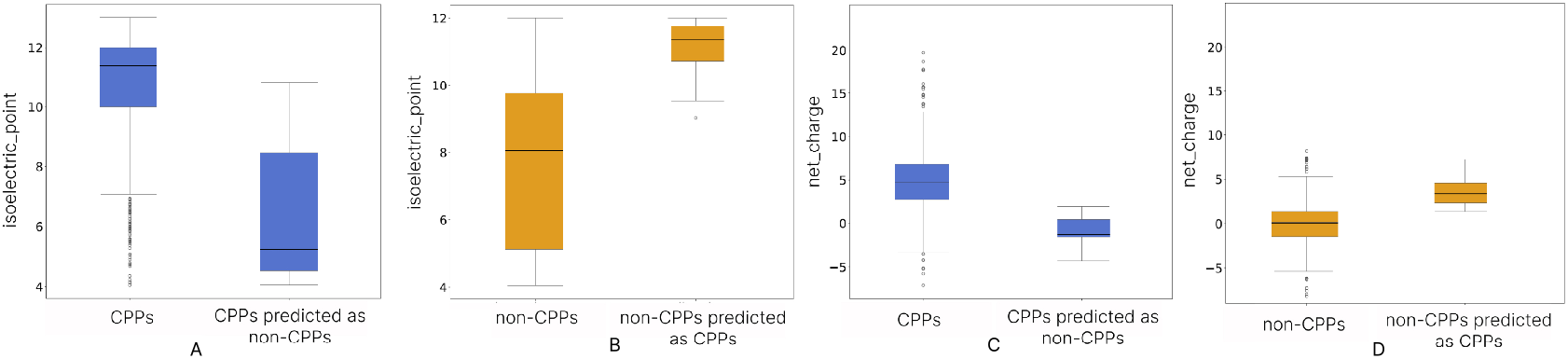
Isoelectric Point and Net Charge Comparison of Train and Misclassified Sequences. **A** the distribution of isoelectric points for positive train sequences and misclassified sequences. **B** isoelectric point distribution for negative train sequences and misclassified sequences. **C** displays the net charge distribution for positive train sequences and misclassified sequences. **D** represents the net charge distribution for negative train sequences and misclassified sequences.

While machine learning models may have a tendency to be more sensitive to false positives or false negatives, our model outperformed all the competitors considering the benchmark datasets used in the literature to validate their results. We had instances that were incorrectly predicted in MLCPP2.0 and CPP1708 independent dataset, meaning sequences that should have been classified as negative but were classified as positive. However, this is not necessarily an error due to the nature of the negative sequences. Given the lack of data, many of these sequences were generated automatically based on experimentally validated negative sequences, so, although unlikely, these instances could potentially be real positive sequences that have not yet been experimentally determined. In this case, our method would not be making an incorrect prediction, but rather identifying a CPP that has not yet been determined experimentally.

### CPP efficiency classifier

The second stage of PERSEUcpp was responsible for assessing the efficiency of peptide penetration. In other words, beyond categorizing a peptide as CPP or non-CPP, the proposed method evaluated how efficient the peptide is in terms of cell penetration.

#### PERSEUcpp compared with the state-of-the-art efficiency predictor

To build the PERSEUcpp efficiency predictor we used the atomic composition (the number of carbon, hydrogen, nitrogen, oxygen, and sulfur atoms normalized by chain length). For our method, the atomic composition played an important role in improving results. The MLCPP2.0 efficiency predictor, then considered the state-of-the-art method, focused on descriptors Composition of K-spaced Amino Acid Group Pairs (CKSAAGP), Dipeptide Composition (DPC), and Amino Acid Composition (AAC).

PERSEUcpp was successful when predicting CPP efficiency, showing compatible or superior results in comparison with the state-of-the-art competitor, as presented in Table 5.

**Table 5.** Comparison results of the proposed PERSEUcpp with MLCPP2.0 balanced dataset. Atm represents the atomic composition group of features.

| Method | MCC | ACC | SN | SP | AUC |
| --- | --- | --- | --- | --- | --- |
| MLCPP2.0 | 0.354 | 0.677 | 0.652 | **0.750** | **0.708** |
| **PERSEUcpp-Atm** | **0.356** | **0.726** | **0.761** | 0.625 | 0.690 |

The bold values denote the best performance among state-of-the-art methods.

#### Feature importance

Since we are dealing exclusively with CPPs and attempting to distinguish them by high and low efficiency, a variety of experiments were conducted to evaluate which feature group is capable of improving classification. The results of these experiments are available in the Supplementary Section 5 in Tables S14-S15. The set of features selected to train the model for determining the efficiency of a CPP were the atomic frequencies, as they presented the best results. Figure S9 in Supplementary Subsection 5.1 provides the detailed importance of each atomic feature.

An important aspect of our model performance is its superior metrics in sensitivity (SN) and accuracy (ACC), and AUC comparable with MLCPP2.0. Our model achieved a SN 0.761, representing a 16.72% increase compared to MLCPP2.0 SN, which is 0.652. SN, which measures the model ability to correctly predict efficient CPPs instances in the context of all the existing positive instances, is a relevant metric for our model as it ensures more accurate identification of efficient CPPs. Additionally, our model presents AUC 0.690, which is close to the value reached by MLCPP2.0. These improvements are important as they indicate our model better capability to correctly identify efficient CPPs, thereby having potential to contribute to the development of more effective therapeutic agents.

## Conclusion

Our study introduces PERSEUcpp, a machine-learning strategy specifically designed for the prediction of cell-penetrating peptides. By leveraging a comprehensive set of descriptors, encompassing physicochemical properties, atomic characteristics, and sequencial features our model, employing the ERT algorithm, demonstrates superior predictive capabilities.

The thoroughness of our method, reflected in the comprehensive nature of the descriptors, not only provides significant biological insights but also outperforms the state-of-the-art methods across all classification metrics on the MLCPP2.0, CPP924, CPP1708 test datasets, which are the independent datasets considered in the literature for evaluating CPP prediction.

Additionally, there is a second stage of PERSEUcpp, which was responsible for assessing the efficiency of peptide penetration, classifying CPPs as low or high efficiency. In this stage, our method presented results comparable or superior to the state-of-the-art.

Our method represents a significant advancement in the computational prediction of CPPs, offering a powerful, reliable and interpretable tool for researchers and practitioners in the field. We anticipate that PERSEUcpp can contribute to the development of drug delivery systems and peptide based therapeutics.

## Data availability

All the data and source code are available at the GitHub repository: https://github.com/goalmeida05/PERSEU

## Acknowledgments

Coordenação de Aperfeiçoamento de Pessoal de Nível Superior – Brasil (CAPES); Conselho Nacional de Desenvolvimento Científico e Tecnológico (CNPq); Fundação de Amparo a` Pesquisa do Estado de Minas Gerais (FAPEMIG).

## 1 Introduction

The CellPPD [1] platform combines ML methods and experimental data to predict CPPs. It focuses on peptide characteristics, including charge, hydrophobicity, and sequence length. Users can submit peptide sequences to the platform, which provides a predictive score indicating the likelihood of cell-penetrating capability based on these features.

CPPpred [2] employs a supervised learning approach to predict the probability of peptides being CPPs. The model uses amino acid composition, physicochemical properties such as molecular weight, isoelectric point, hydrophobicity, and specific sequence motifs commonly found in CPPs. By calculating a score for each peptide, CPPpred allows threshold adjustments to manage false positives.

BChemRF [3], is a ML model for CPP prediction. Using a Random Forest classifier, BChemRF extracts features from peptide sequences, including physicochemical properties, amino acid composition and arrangement, predicted secondary structures, flexibility, and evolutionary information. The ensemble learning approach of BChemRF enhances prediction accuracy and reliability by capturing complex patterns associated with cell-penetrating capability.

SkipCPP-Pred [4] uses a k-skip-n-gram model to convert variable sequence lengths into fixed-length feature vectors. This approach captures the frequency of n residues separated by up to k other residues, including both contiguous and non-contiguous pairs to retain spatial information. The adaptive feature representation adjusts to the length of each peptide, ensuring flexible and inclusive feature extraction. These features are then used to train a ML model to predict the cell-penetrating potential of peptides.

MLCPP2.0 [5], a two-layer classifier named MLCPP2.0 was proposed to predict CPPs and their uptake efficiency. The model uses SVM to predict CPPs based on various features. This predictor achieved superior performance compared to existing predictors and identified independent features for CPP classification. The features used include dipeptide and amino acid compositions, frequency and transition patterns, binary representations, substitution matrices, and physicochemical properties. These groups of features are employed to enhance the accuracy of CPP prediction. The major limitation of this model is that false negatives predicted at the first layer will not be processed by the second layer.

The machine learning model implemented in MLCPP2.0 [5] for predicting the uptake efficiency of cell-penetrating peptides (CPPs) is a meta-model constructed using 11 groups of feature sets. The selected model was the Gradient Boosting with the top 50 feature set. The features utilized included various peptide characteristic encodings such as Amino Acid Composition (AAC), Quasi-Sequence Order (QSO), Composition, Transition, and Distribution (CTDC), Dipeptide Deviation from Expected Mean (DDE), Composition of K-Spaced Amino Acid Pairs (CKS), Dipeptide Composition (DPC), and Composition of K-Spaced Amino Acid Groups Pair (CKSAAGP), among others. These encodings were determined to be important through SHapley Additive exPlanations (SHAP) analysis, a method that measures the contribution of each feature to the model’s predictions based on Shapley values from game theory [6], indicating their role in predicting CPPs in the second layer model.

Recently, the SiameseCPP [7] predictor demonstrated that contrastive learning can extract intrinsic characteristics of CPPs. SiameseCPP is based on deep neural networks, specifically a Siamese network, which learns discriminative features directly from peptide sequences. This method automatically extracts relevant patterns and motifs from the sequences without predefined features, generating high-dimensional representations that capture essential characteristics of the peptides. SiameseCPP improves the accuracy of CPP prediction by identifying complex patterns within the sequences. SiameseCPP achieves superior performance compared to MLCPP2.0, but the study still faces some limitations: of note, lacks interpretability, failing to provide practical biological insights.

PractiCPP [8] is a deep learning framework designed for CPP prediction in highly imbalanced datasets, addressing real-world challenges where positive samples are scarce. The model integrates hard negative sampling to refine decision boundaries and employs three feature types: sequential features from amino acid sequences, local features derived from Morgan fingerprints (a molecular descriptor capturing the local structural environment of atoms [9]), and pretrained embeddings from the ESM-2 language model. PractiCPP outperforms state-of-the-art models like SiameseCPP and MLCPP2.0 on both balanced and imbalanced datasets, achieving superior metrics such as AUPR and F1 score.

GraphCPP [10] leverages graph neural networks (GNNs) to predict cell-penetrating peptides (CPPs) by modeling peptide sequences as molecular graphs. This method captures intricate relationships between amino acids and their physicochemical properties, outperforming traditional ML models and sequence-based predictors. By integrating node and edge features, GraphCPP provides robust embeddings that enhance the accuracy of CPP prediction. Its superior performance on balanced and imbalanced datasets establishes GraphCPP as a state-of-the-art tool for CPP identification, addressing challenges such as sequence variability and complex molecular interactions.

## 2 Materials and methods

### 2.1 Feature engineering

**Figure S1:**
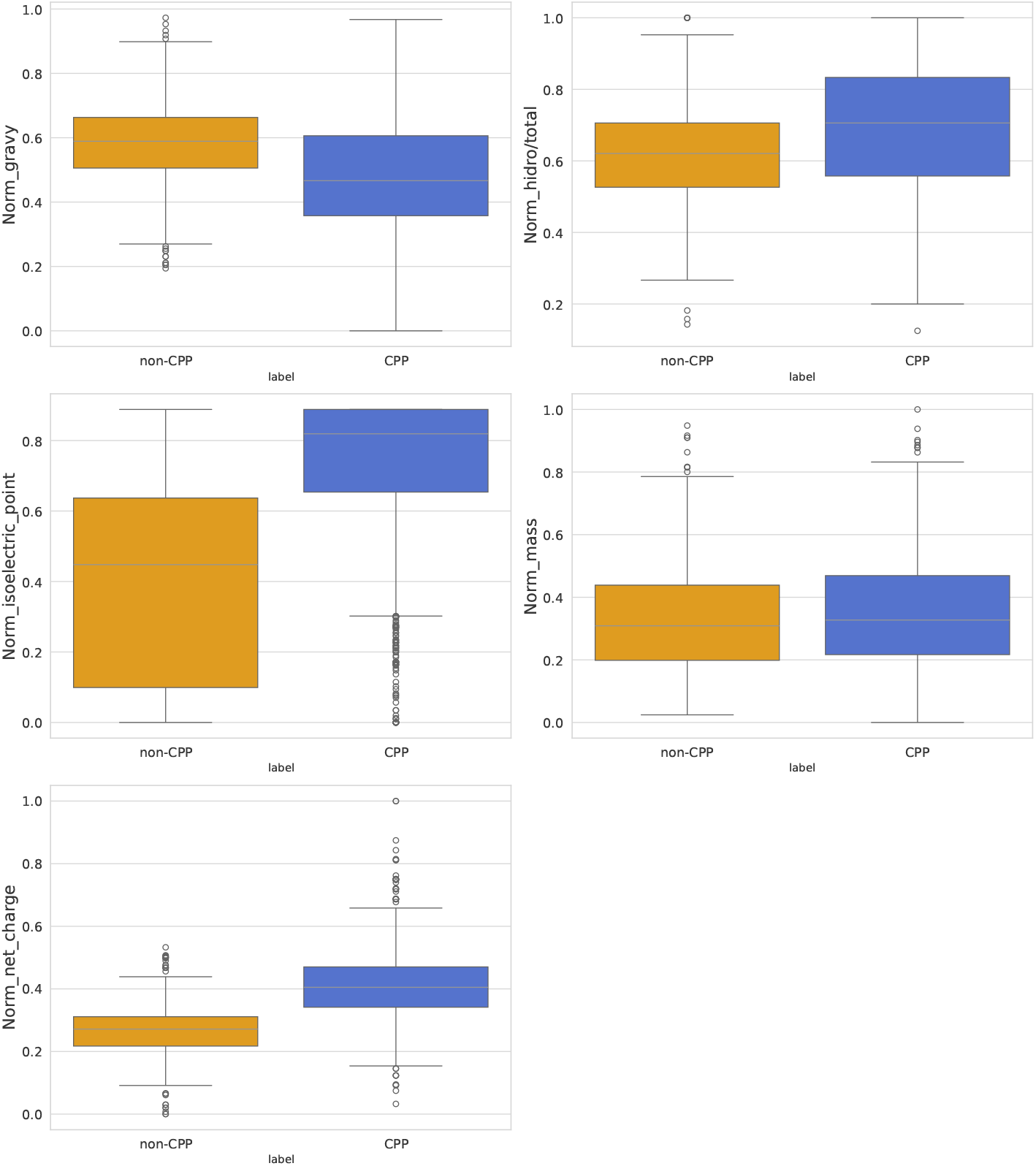
Boxplot of CPPs and Non-CPPs based on physicochemical features on training dataset. CPPs exhibit distinct distributions in features such as hydrophobicity and charge, highlighting their physicochemical differences.

**Figure S2:**
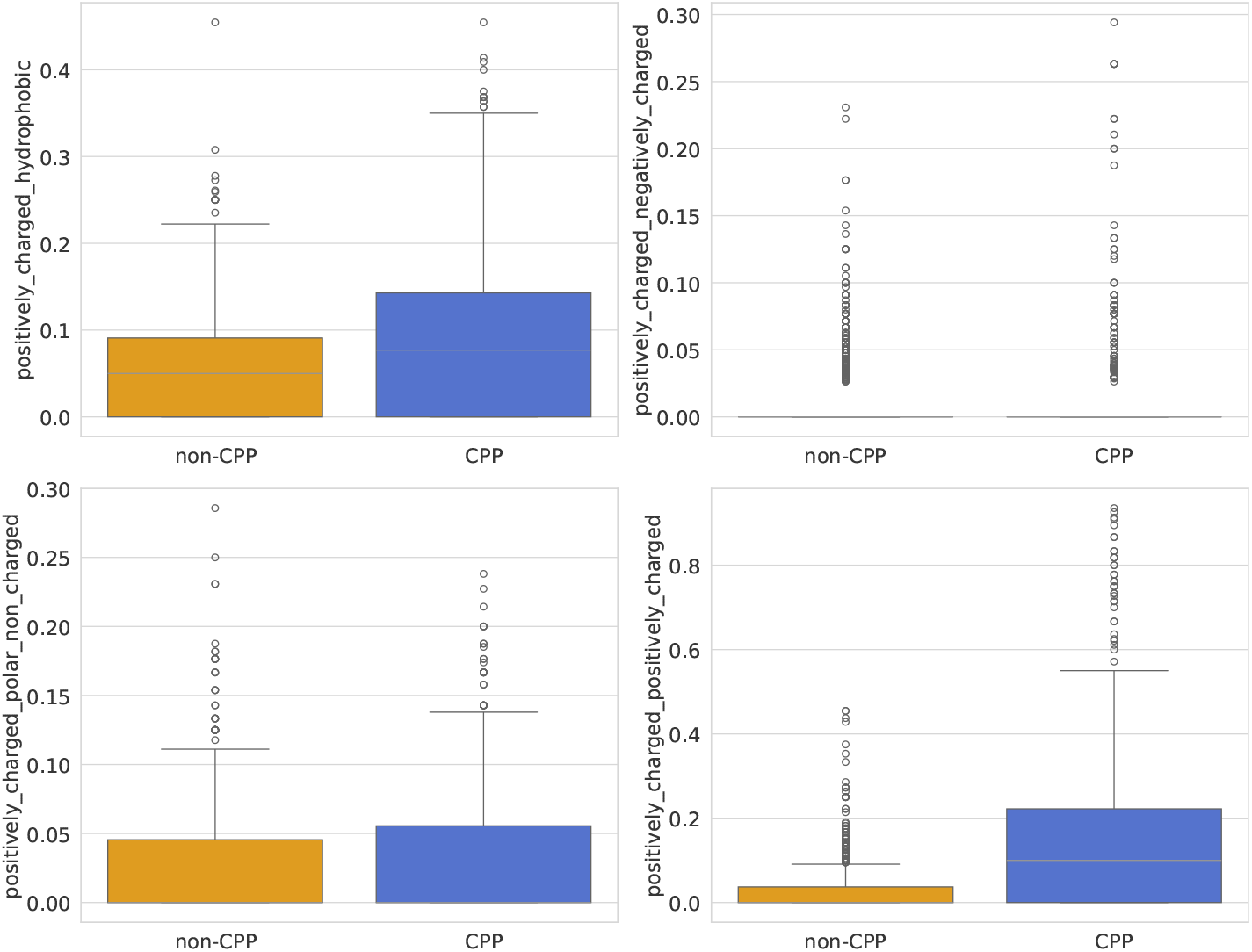
Boxplot of CPPs and Non-CPPs based on CKSAAGP positive charged normalized features on training dataset. CPPs exhibit distinct distributions in features such as positively_charged_hydrophobic, positively_charged_negatively_charged, positively_charged_polar_non_charged and positively_charged_positively_charged, highlighting their CKSAAGP differences.

**Figure S3:**
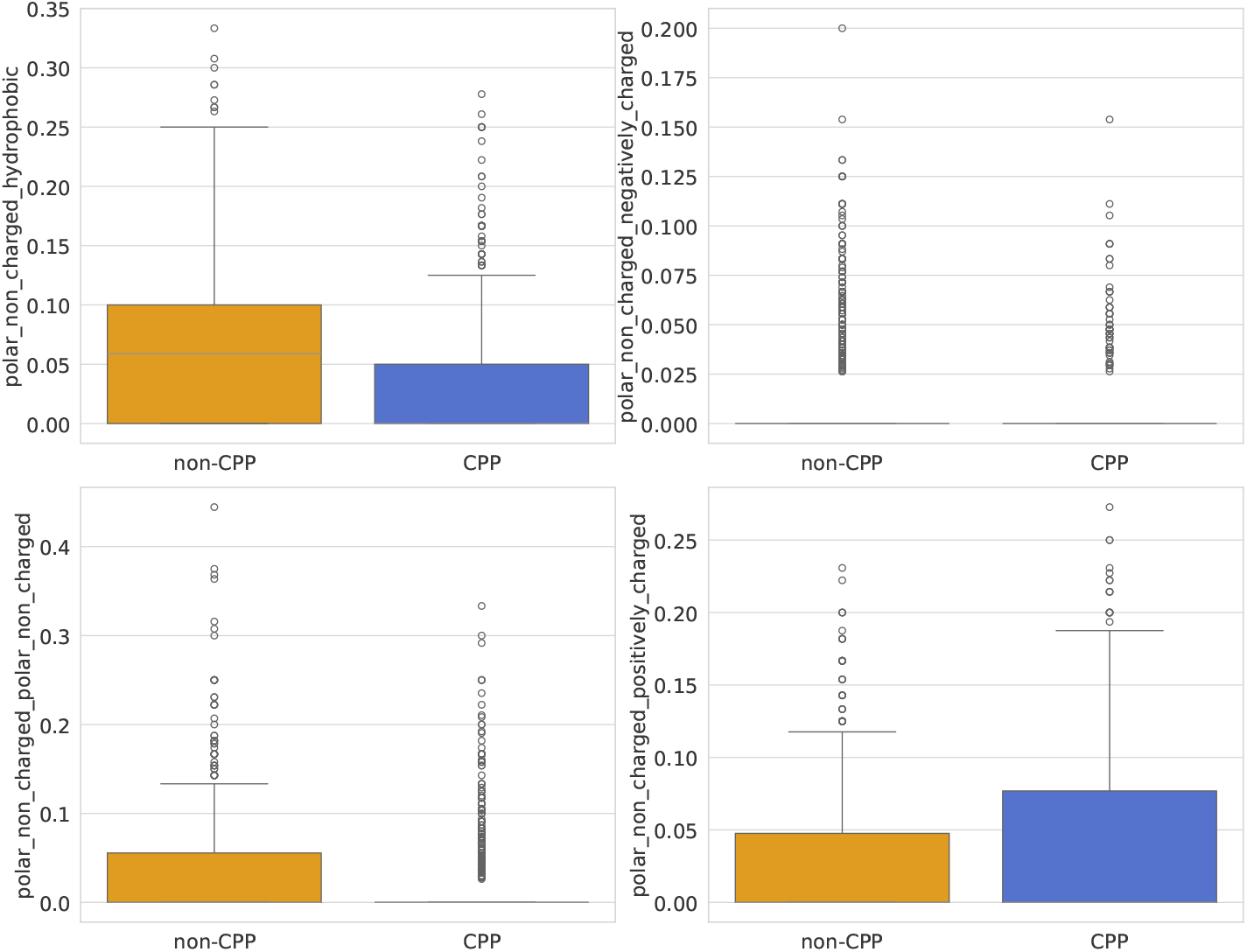
Boxplot of CPPs and Non-CPPs based on CKSAAGP polar non charged normalized features on training dataset. CPPs exhibit distinct distributions in features such as polar_non_charged_hydrophobic, polar_non_charged_negatively_charged, polar_non_charged_polar_non_charged and polar_non_charged_positively_charged, highlighting their CKSAAGP differences.

**Figure S4:**
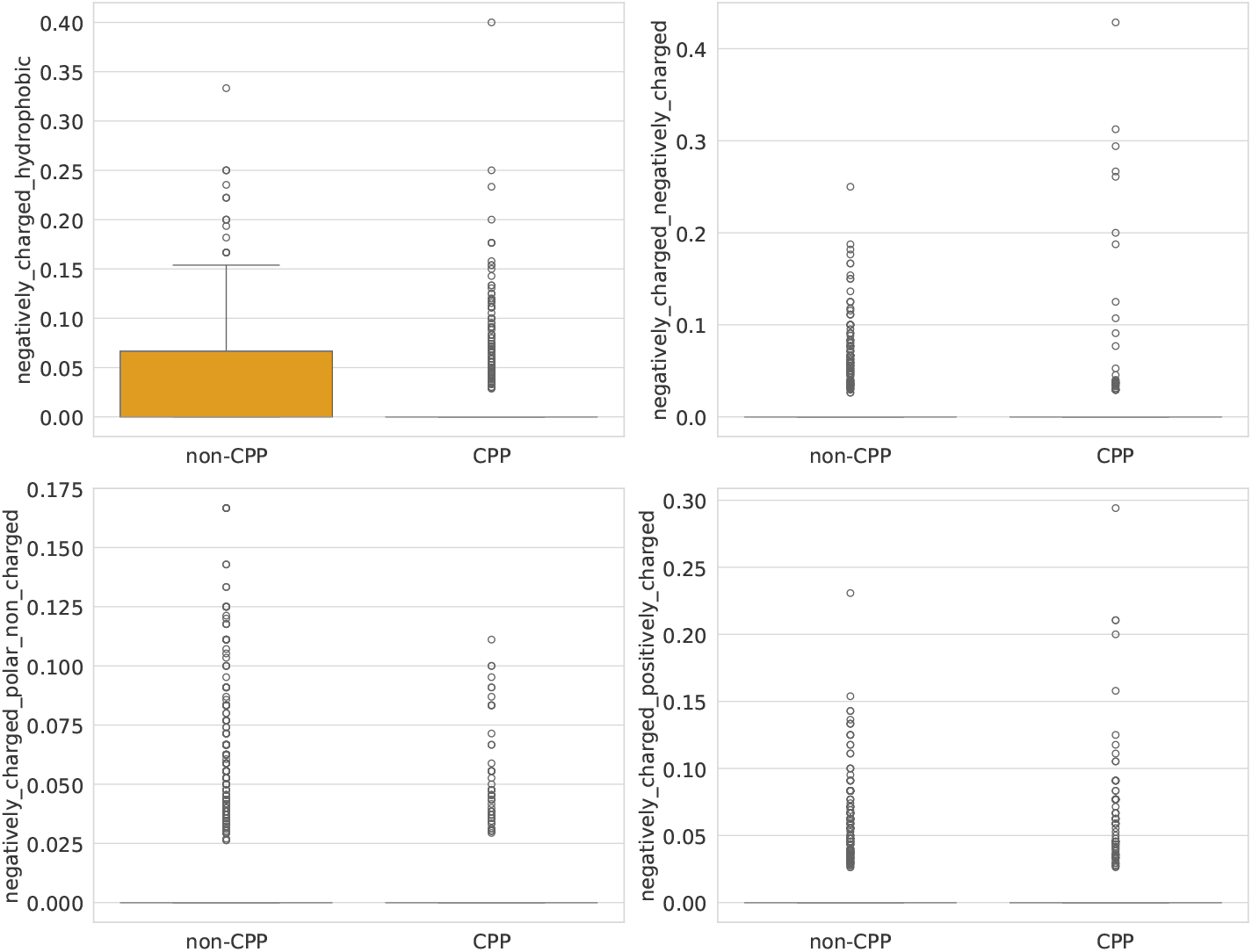
Boxplot of CPPs and Non-CPPs based on CKSAAGP negatively charged normalized features on training dataset. CPPs exhibit distinct distributions in features such as negatively_charged_hydrophobic, negatively_charged_negatively_charged, negatively_charged_polar_non_charged and polar_non_charged_positively_charged, highlighting their CKSAAGP differences.

**Figure S5:**
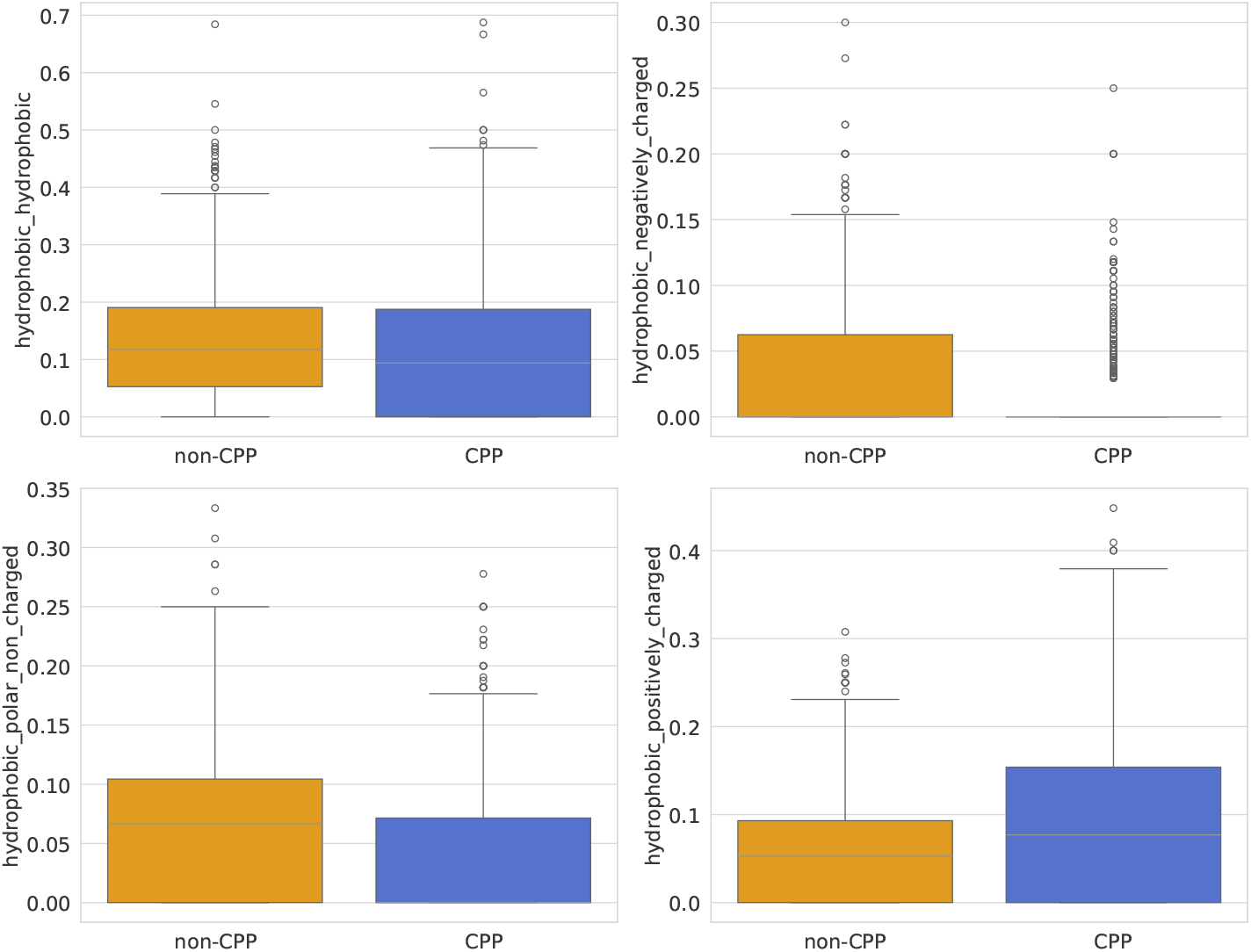
Boxplot of CPPs and Non-CPPs based on physicochemical features on training dataset. CPPs exhibit distinct distributions in features such as hydrophobicity and charge, highlighting their physicochemical differences.

**Figure S6:**
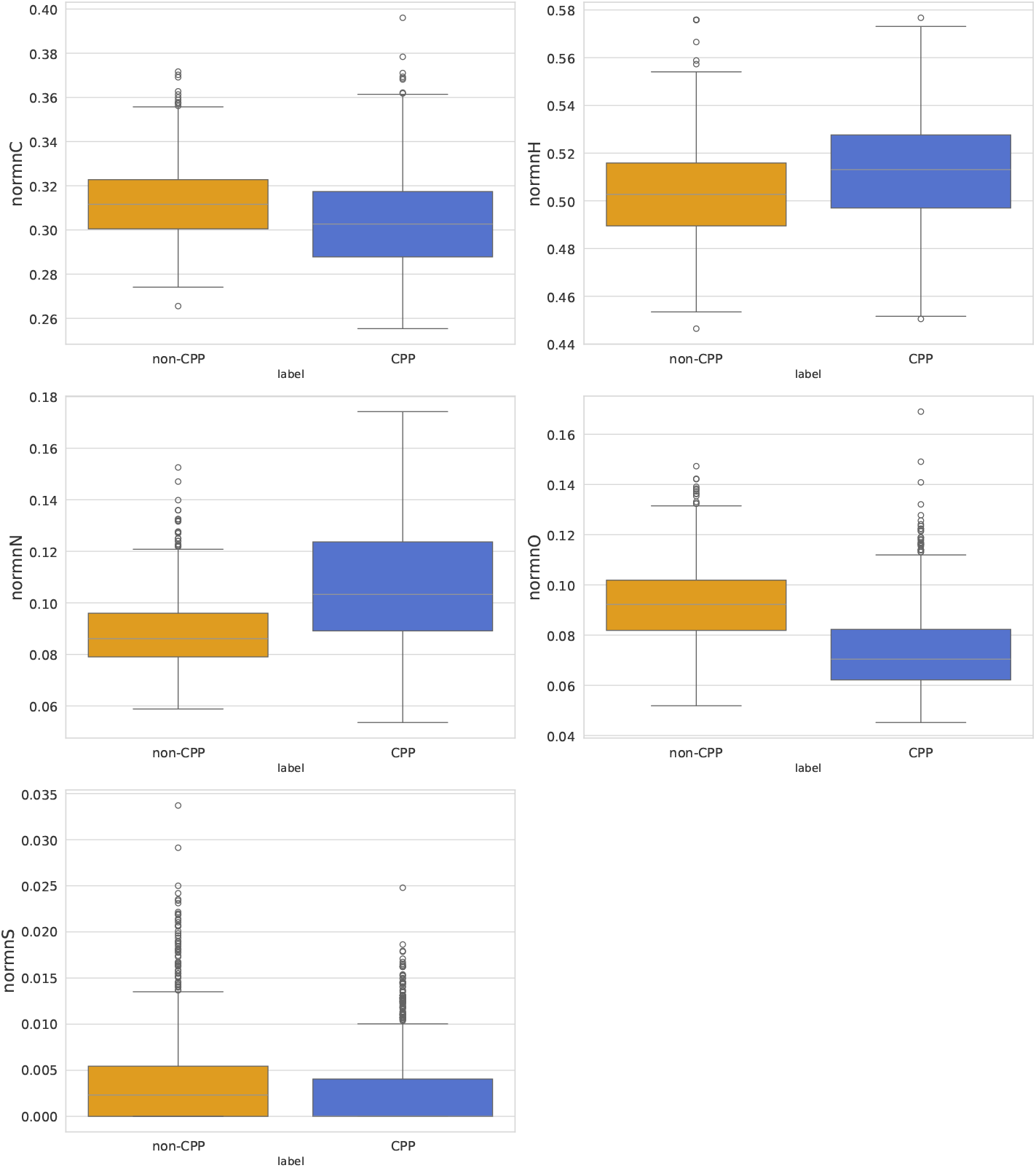
Boxplot of CPPs and Non-CPPs based on physicochemical features on training dataset. CPPs exhibit distinct distributions in features such as hydrophobicity and charge, highlighting their physicochemical differences.

### 2.2 Comparison with alternative methods

Sensitivity, Equation 1, measures the fraction of positive instances correctly identified by the classifier, ranging from 0 to 1. Higher SN means fewer false negatives.

Specificity, Equation 2, measures the fraction of negative instances correctly identified, also ranging from 0 to 1, with higher values indicating fewer false positives.

Accuracy, Equation 3, which measures the overall correctness of the model predictions, also ranges from 0 to 1 but may not be adequate for imbalanced datasets.

The MCC, Equation 4, is a robust measure for binary classification, ranging from -1 (inverse predictions) to +1 (perfect predictions).

The AUC plots the true positive rate against the false positive rate, and represents the model ability to distinguish between classes. An AUC of 0.5 suggests no discriminative ability, with increasing values above 0.5 indicating progressively better discriminative performance. With exception of ACC, all the metrics are robust and appropriate even for scenarios with imbalanced datasets. TP (True Positive) represents the number of cases correctly classified as positive, while FN (False Negative) refers to the cases that are incorrectly classified as negative despite being positive. FP (False Positive) indicates the number of cases incorrectly classified as positive when they are actually negative, and TN (True Negative) represents the cases correctly classified as negative

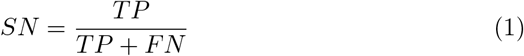

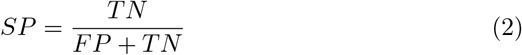

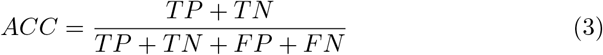

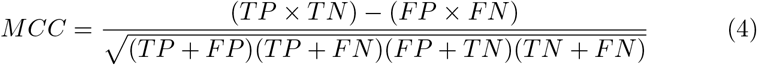

## 3 Results and Discussion

### 3.1 PERSEUcpp cross-validation results

We tested different configurations of the dataset, keeping the same data and features but applying different treatments to each. The datasets were as follows:

- A matrix in which all the 1,137 features are normalized;
- 99 matrices using the dimensionality reduction technique (SVD) on normalized features, varying the number of singular values from 2 to 100, incrementing by 1.

A large number of features can introduce bias into a machine learning model, mainly if some of these features are not relevant to the classification task. This issue can lead to overfitting, a phenomenon where the model learns patterns that are specific to the training set and fails to generalize effectively to new data. To address this, we used feature importance scores from the best-performing algorithms ERT and RF (Table S1) to identify the most relevant features. Based on this analysis, we reduced the feature set from 1,137 to 522 features. This approach ensured that only the most informative features were retained for further modeling.

Table S1 presents the best results obtained by the algorithms, indicating that the EXT with the top 522 normalized features achieved higher performance in the classification of peptides. Therefore, the ERT algorithm was selected for the next steps. The subsection *PERSEUcpp cross-validation results*, in Tables S2-S11contains the performance data of all the experiments conducted.

**Table S1:**
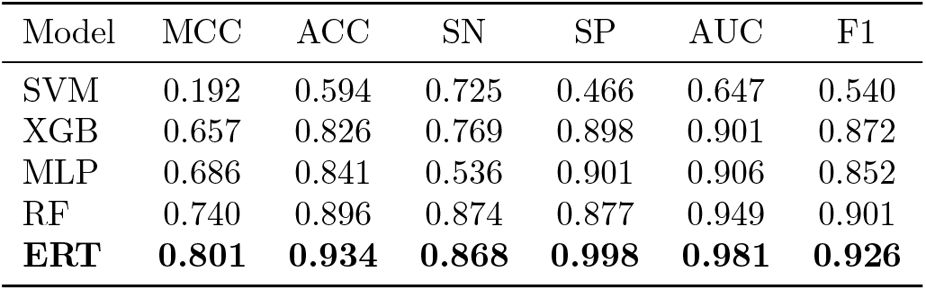
Performance comparison of machine learning algorithms for CPP prediction using the same dataset. The performance of five machine learning models SVM, XGB, MLP, RF, and ERT in predicting cell-penetrating peptides (CPPs).

**Table S2:** Performance classification of Extremelly Randomized Trees in independent datasets.

| Features | MLCPP2.0 Independent Dataset |  |  |  |  |  | CPP924 Independent Datasets |  |  |  |  |  |
| --- | --- | --- | --- | --- | --- | --- | --- | --- | --- | --- | --- | --- |
|  | MCC | ACC | SN | SP | F1-Score | AUC | MCC | ACC | SN | SP | F1-Score | AUC |
| TOP30 | 0.662 | 0.969 | 0.763 | 0.978 | 0.673 | 0.949 | 0.765 | 0.881 | 0.789 | 0.956 | 0.857 | 0.961 |
| TOP90 | 0.778 | 0.981 | 0.817 | 0.989 | 0.787 | 0.968 | 0.839 | 0.917 | 0.828 | 0.989 | 0.900 | 0.989 |
| TOP150 | 0.813 | 0.984 | 0.860 | 0.989 | 0.820 | 0.979 | 0.861 | 0.929 | 0.855 | 0.989 | 0.915 | 0.992 |
| TOP210 | 0.829 | 0.986 | 0.849 | 0.992 | 0.835 | 0.980 | 0.883 | 0.941 | 0.881 | 0.989 | 0.931 | 0.993 |
| TOP270 | 0.839 | 0.987 | 0.860 | 0.989 | 0.836 | 0.979 | 0.906 | 0.952 | 0.907 | 0.989 | 0.945 | 0.994 |
| TOP330 | 0.809 | 0.985 | 0.860 | 0.989 | 0.816 | 0.980 | 0.906 | 0.952 | 0.908 | 0.989 | 0.945 | 0.991 |
| TOP360 | 0.837 | 0.986 | 0.871 | 0.990 | 0.843 | 0.989 | 0.929 | 0.964 | 0.934 | 0.989 | 0.959 | 0.994 |

**Table S3:**
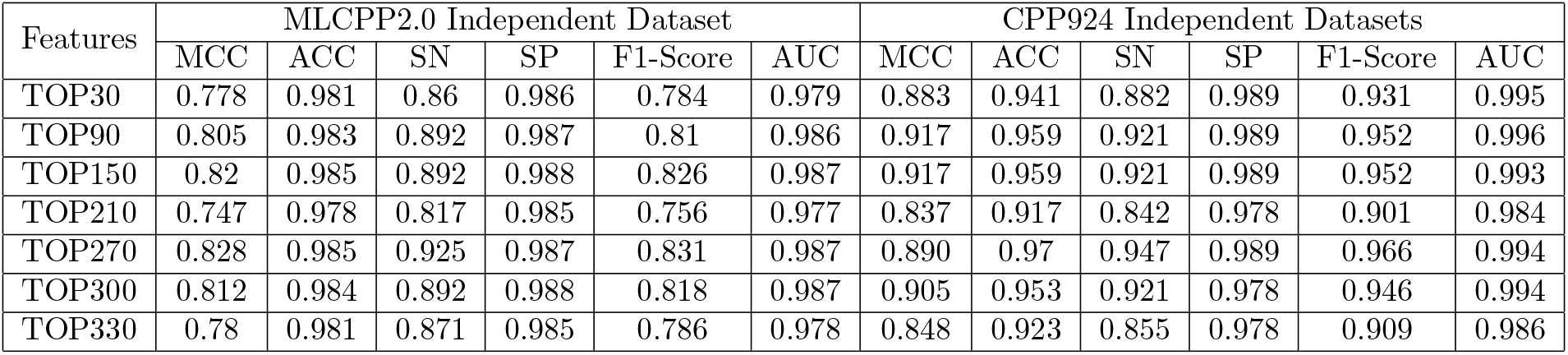
Performance classification of Random Forest Classifier in independent datasets.

**Table S4:**
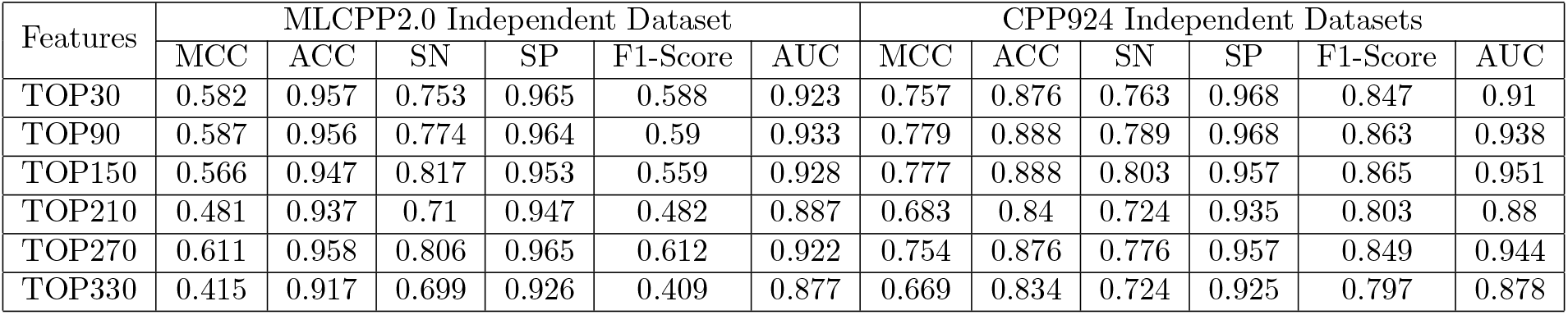
Performance classification of Multi-Layer Perceptron in independent datasets.

**Table S5:**
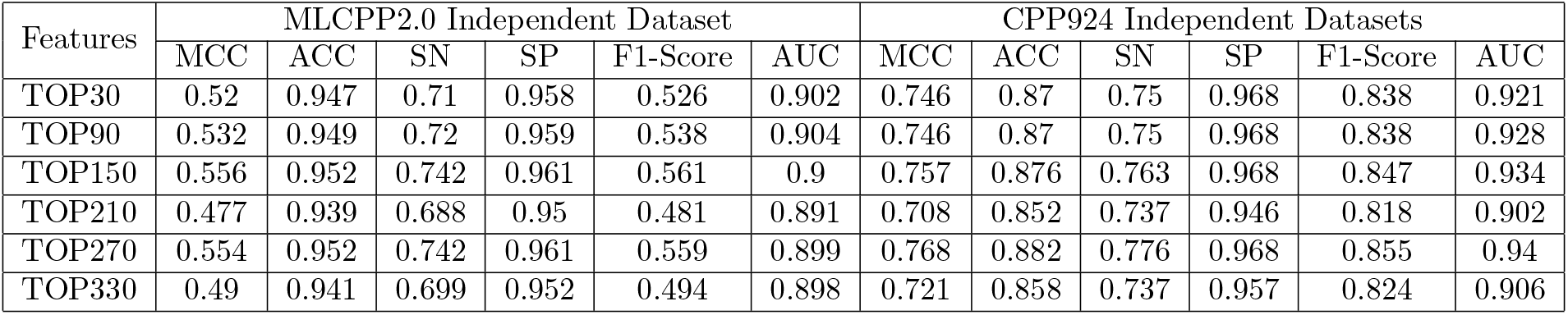
Performance classification of SVM in independent datasets.

**Table S6:**
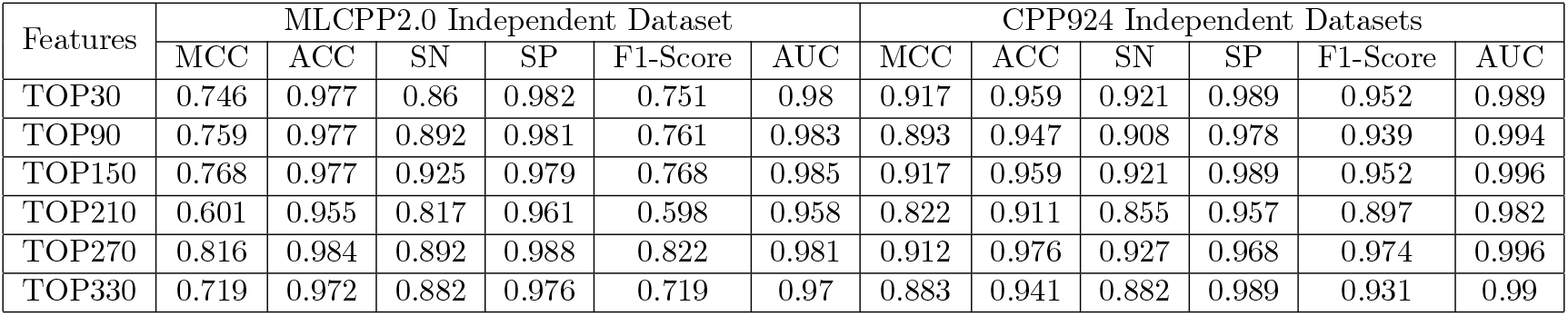
Performance classification of LGBM in independent datasets.

**Table S7:**
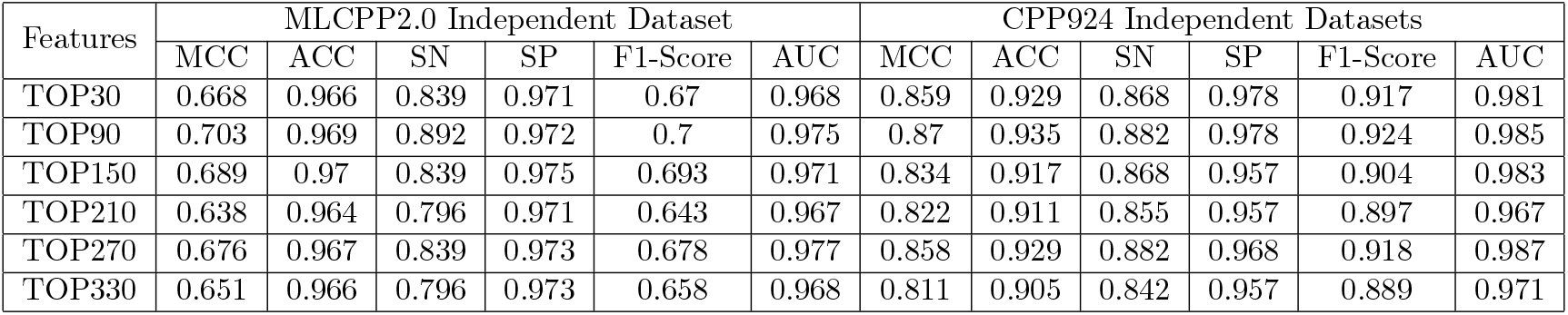
Performance classification of Gradient Boost in independent datasets.

**Table S8:**
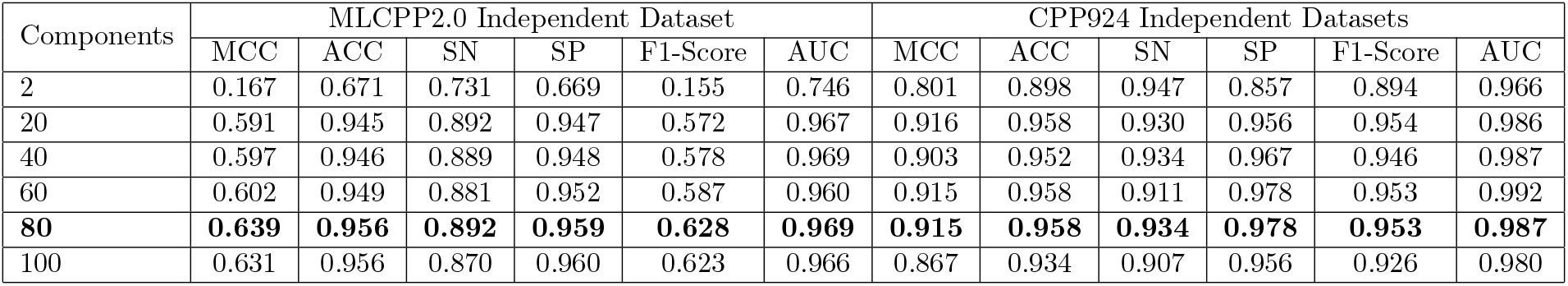
Performance classification with SVD and Extremely Randomized Trees in the independent datasets.

**Table S9:**
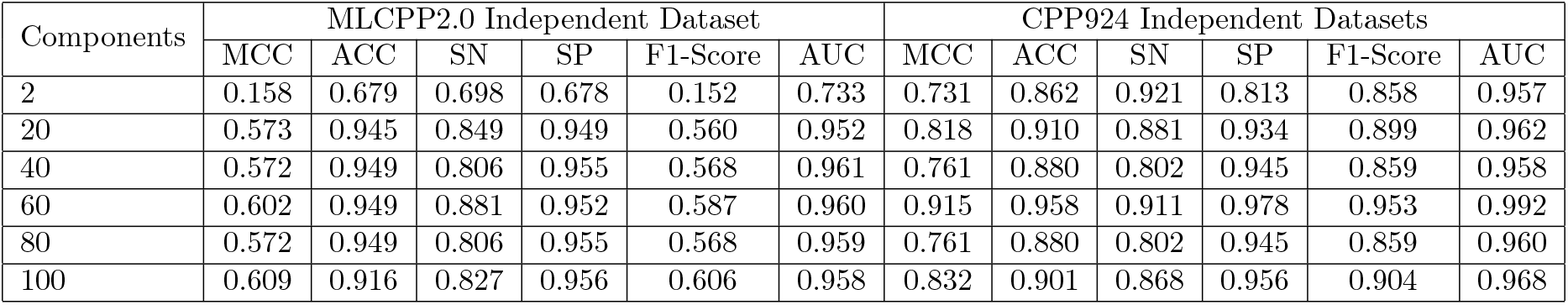
Performance classification with SVD and Random Forest Classifier in the independent datasets.

**Table S10:**
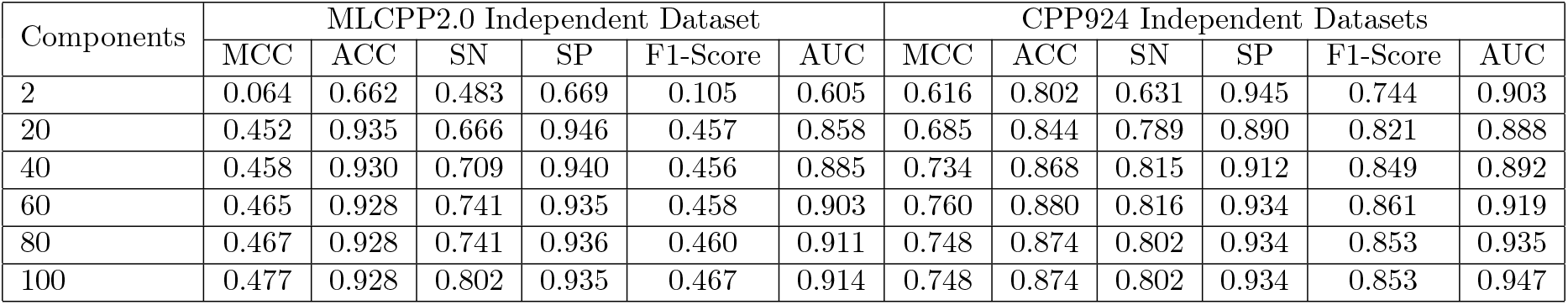
Performance classification with SVD and Logistic Regression in the independent datasets.

**Table S11:**
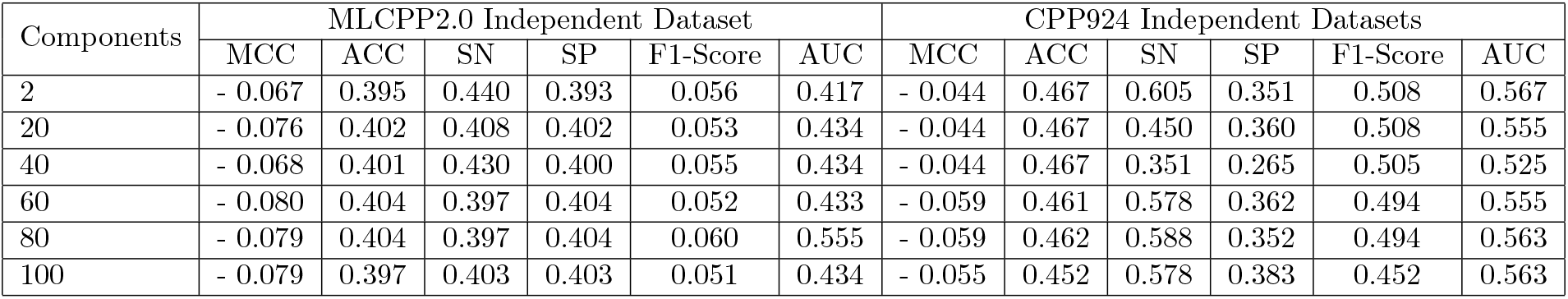
Performance classification with SVD and SVM in the independent datasets.

### 3.2 Feature Importance

**Table S12:**
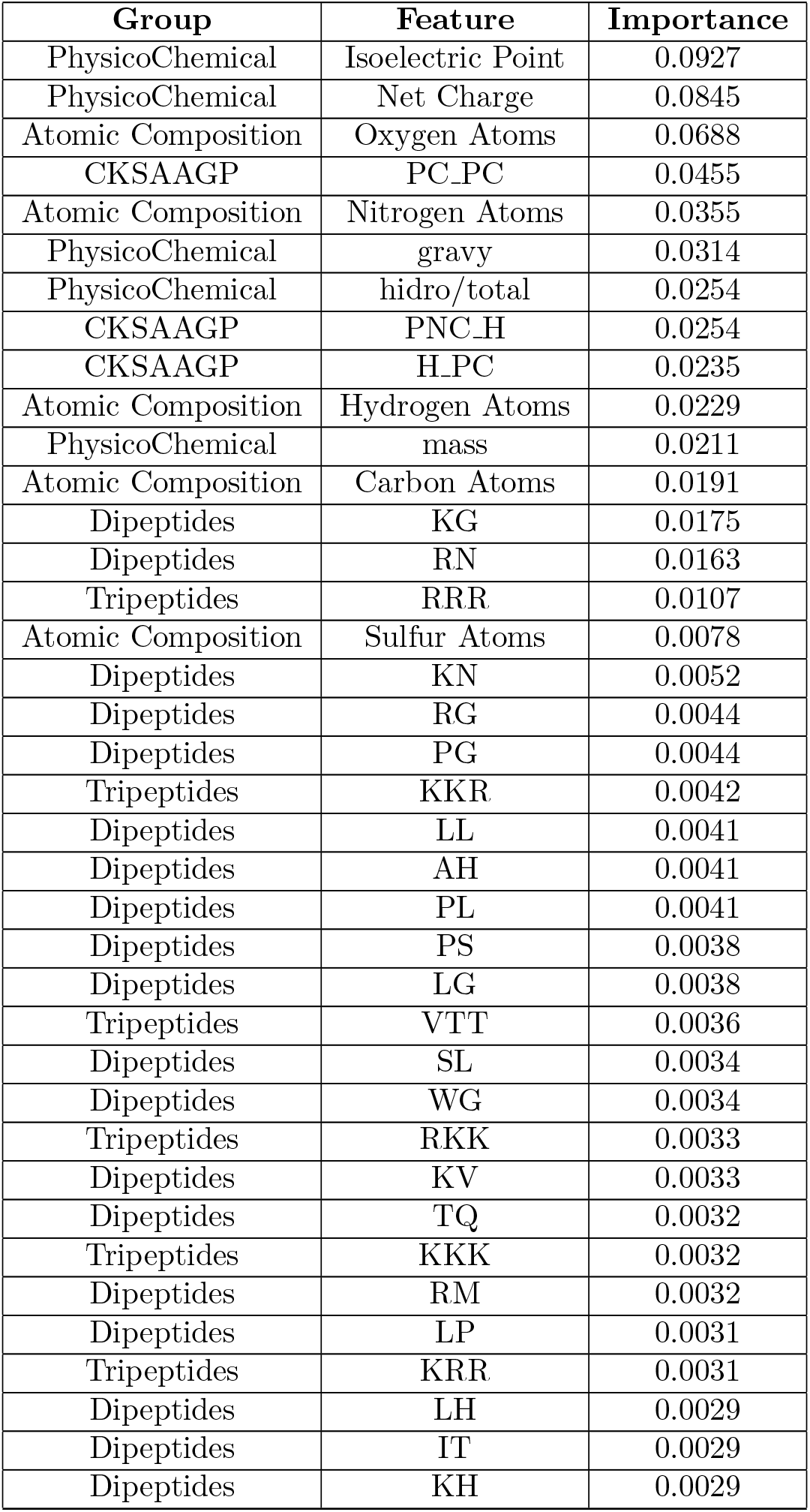
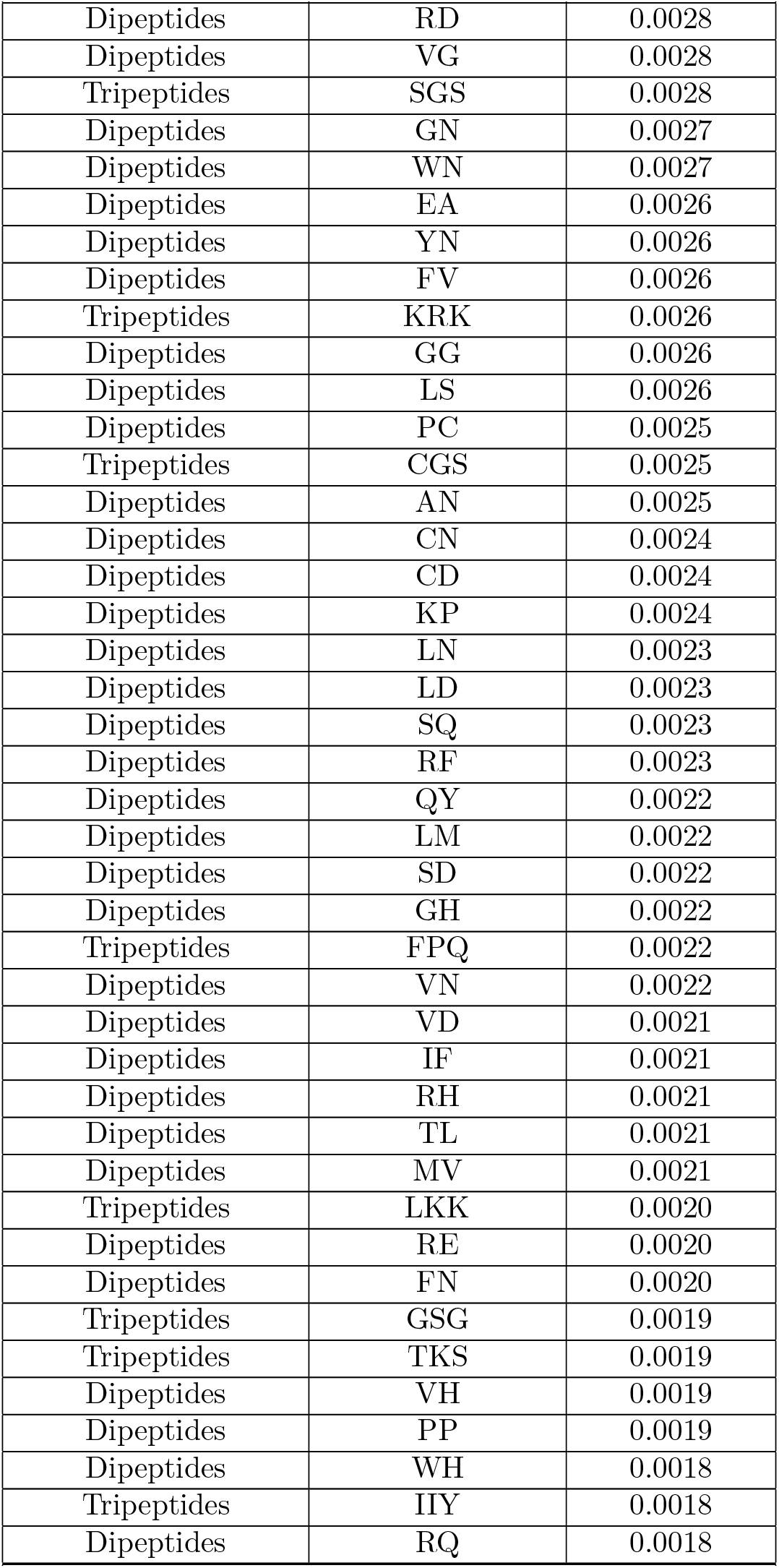
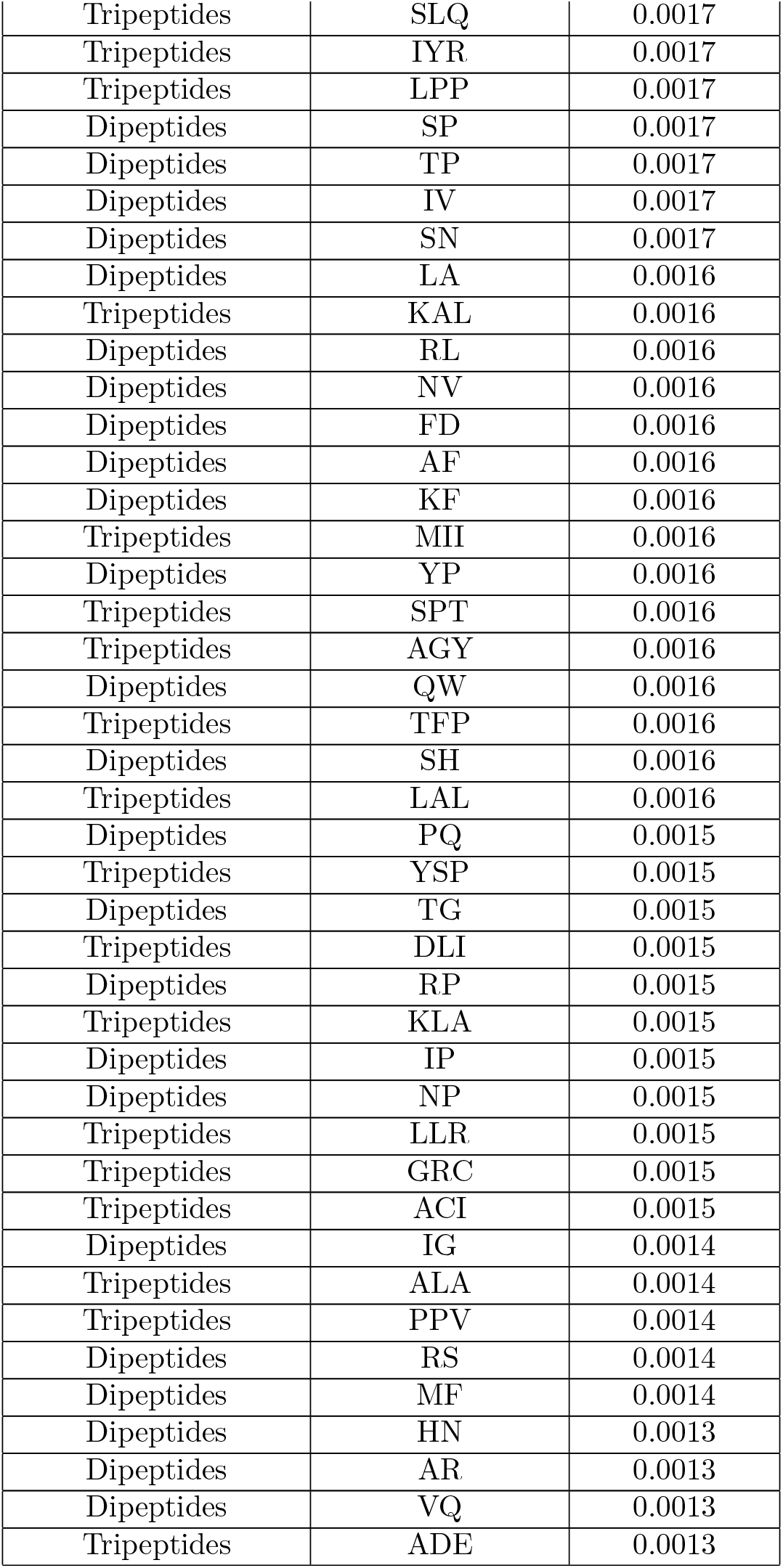
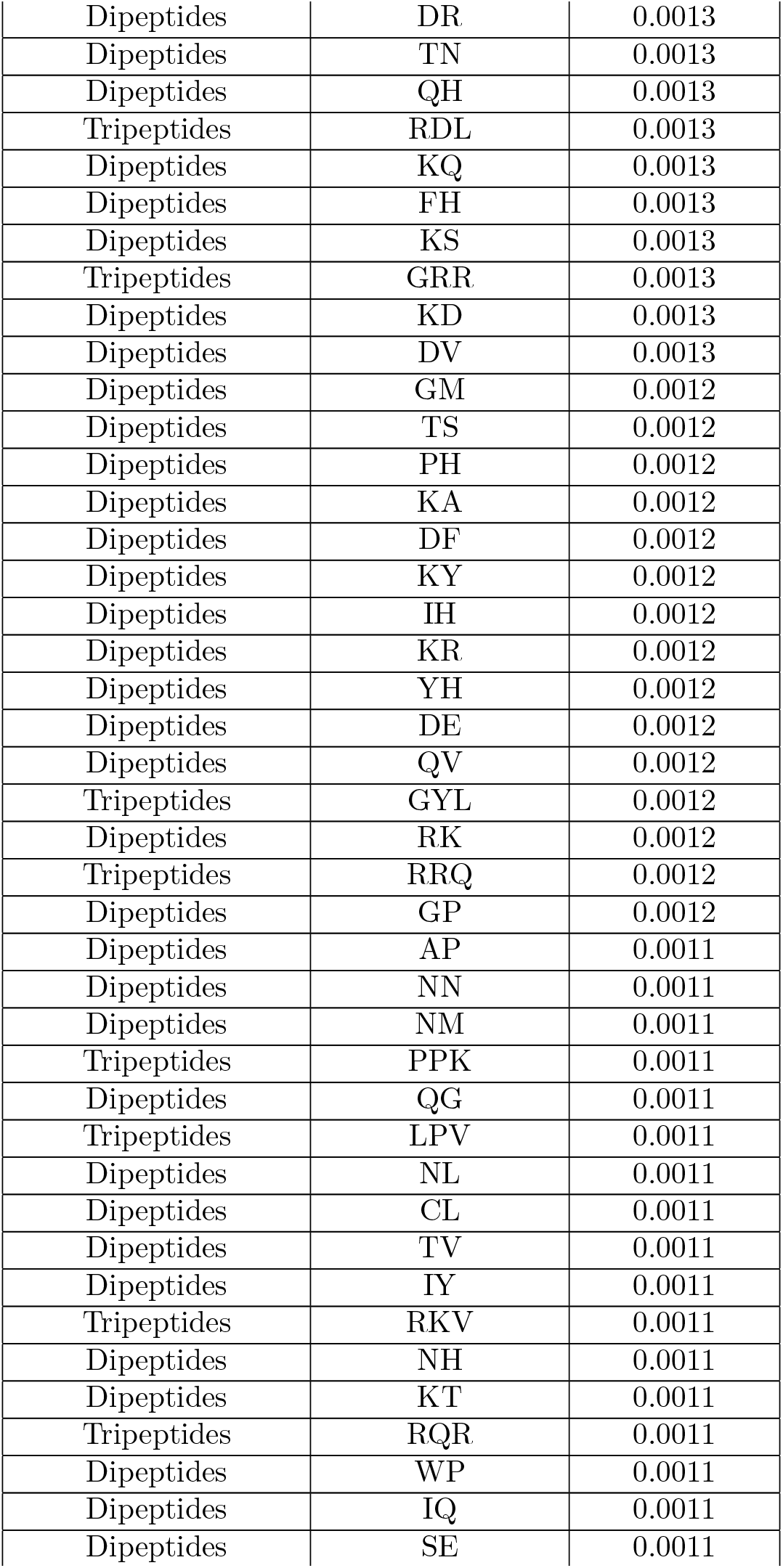
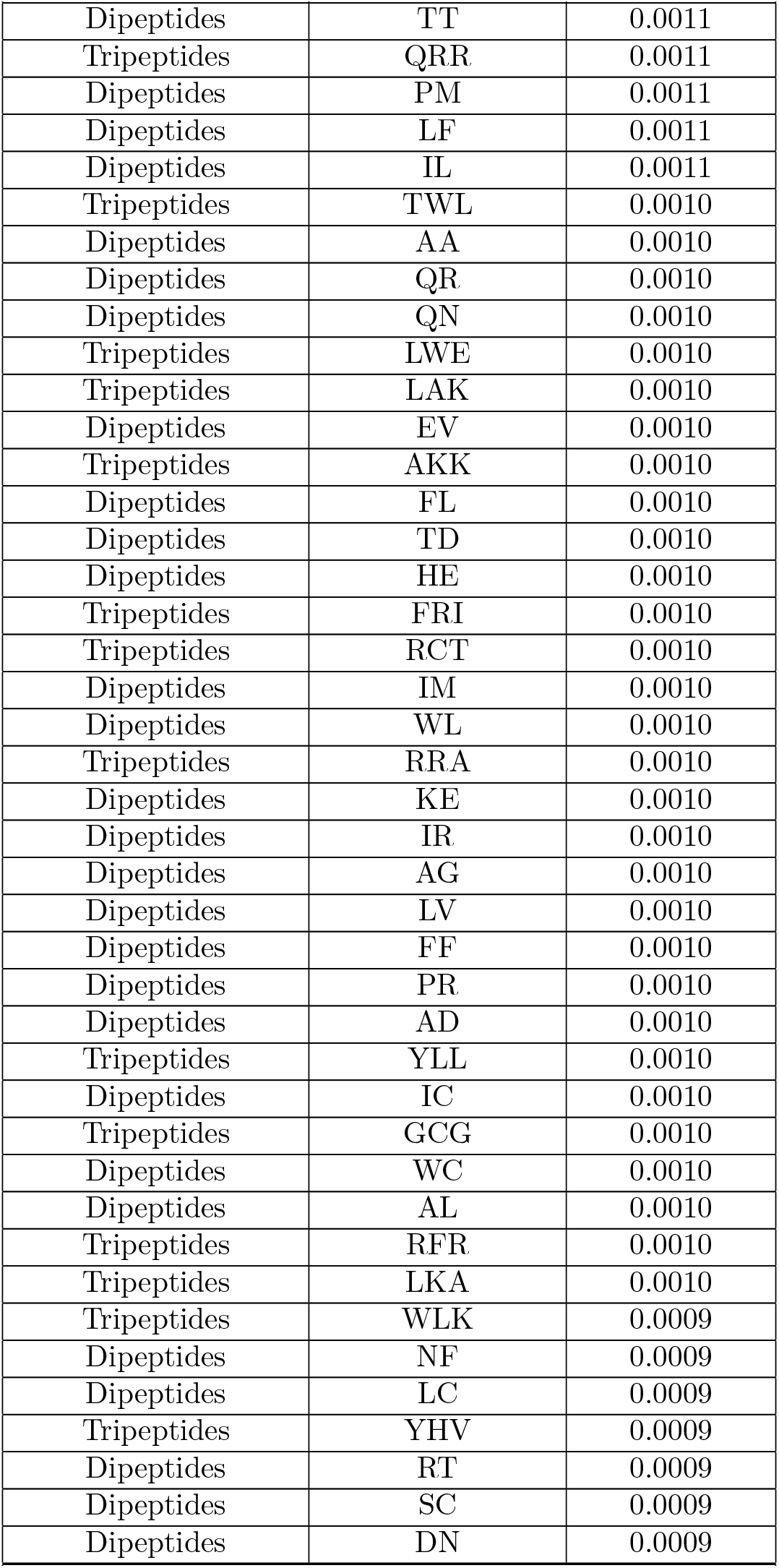
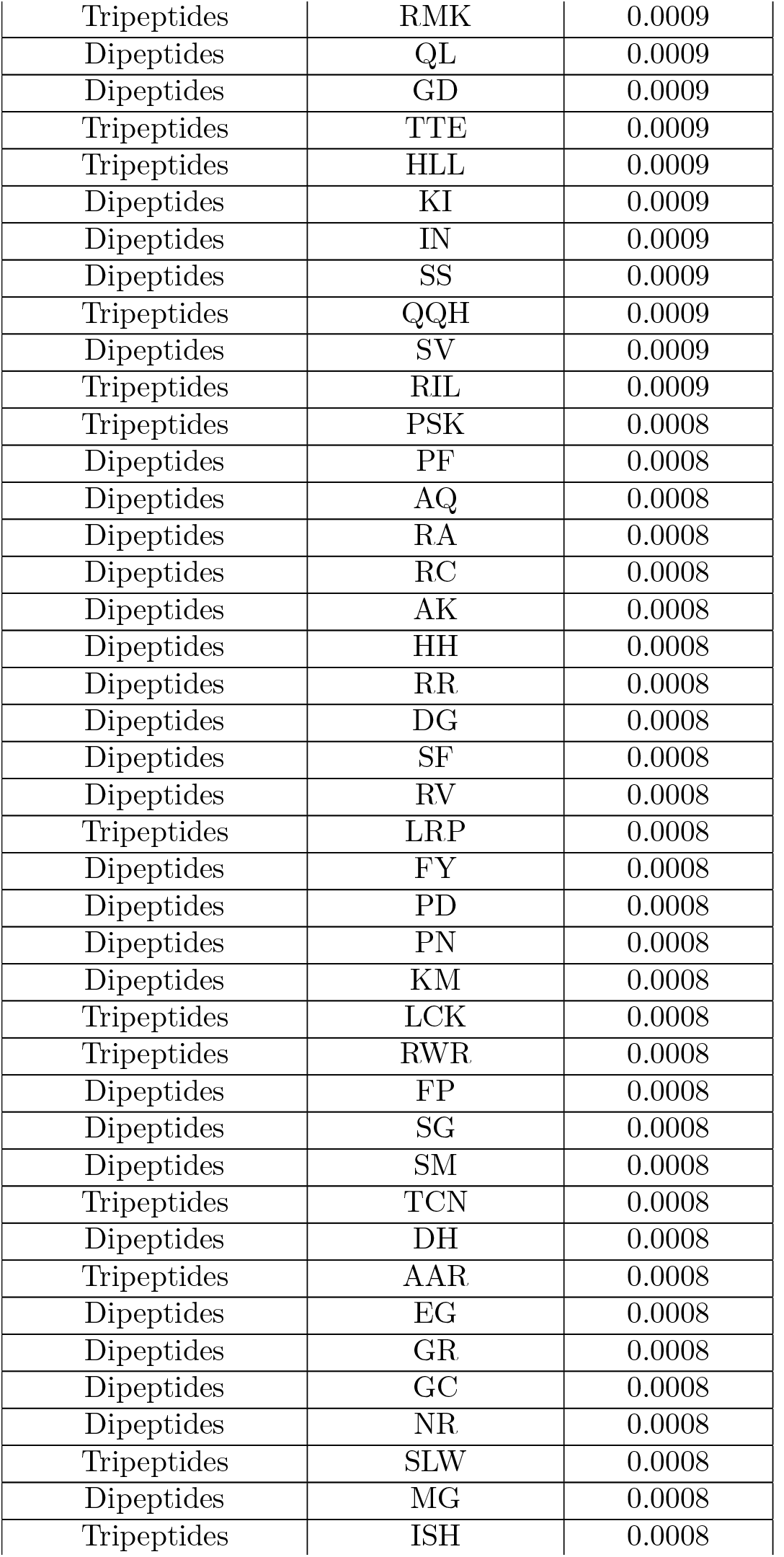
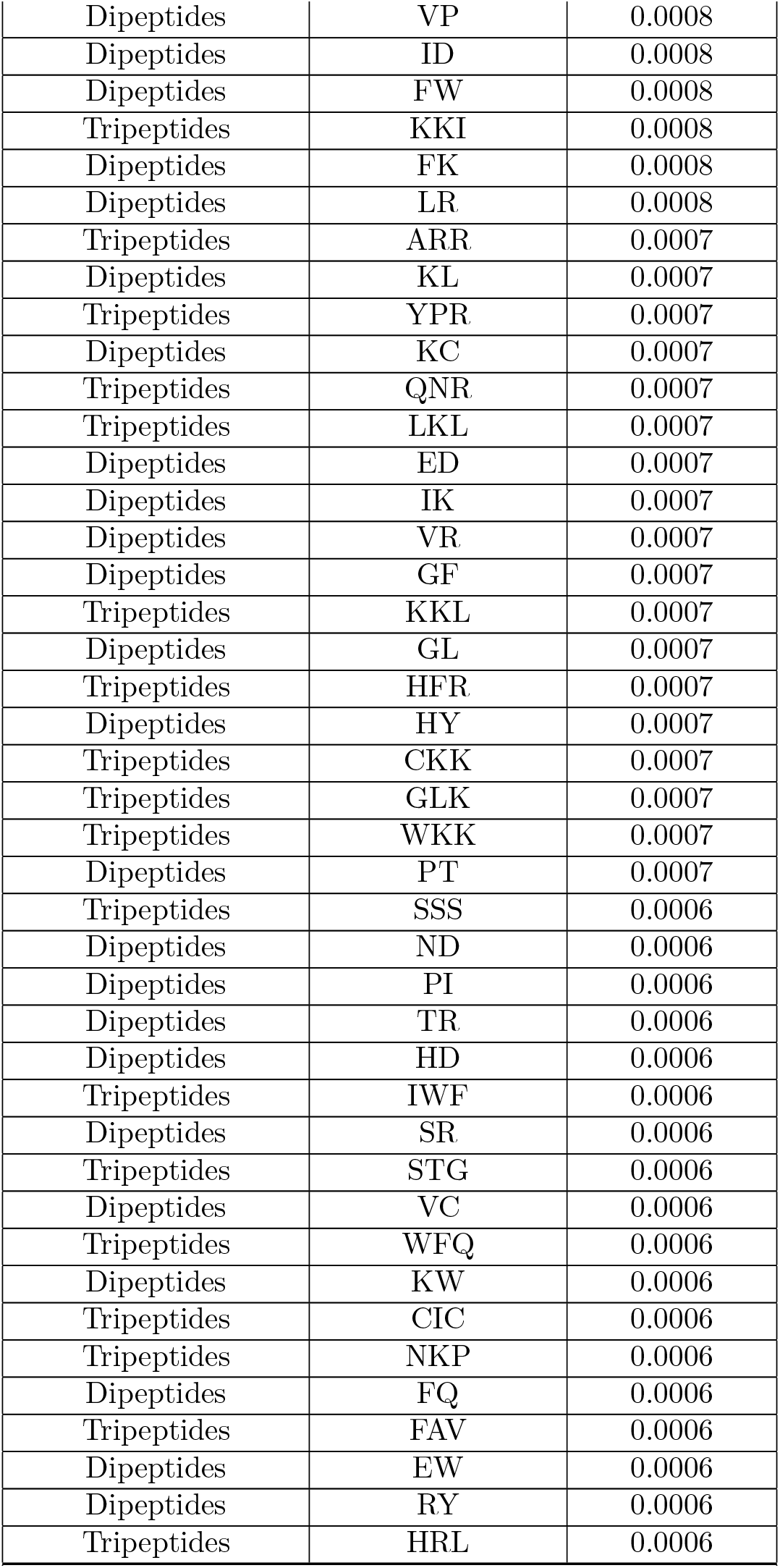
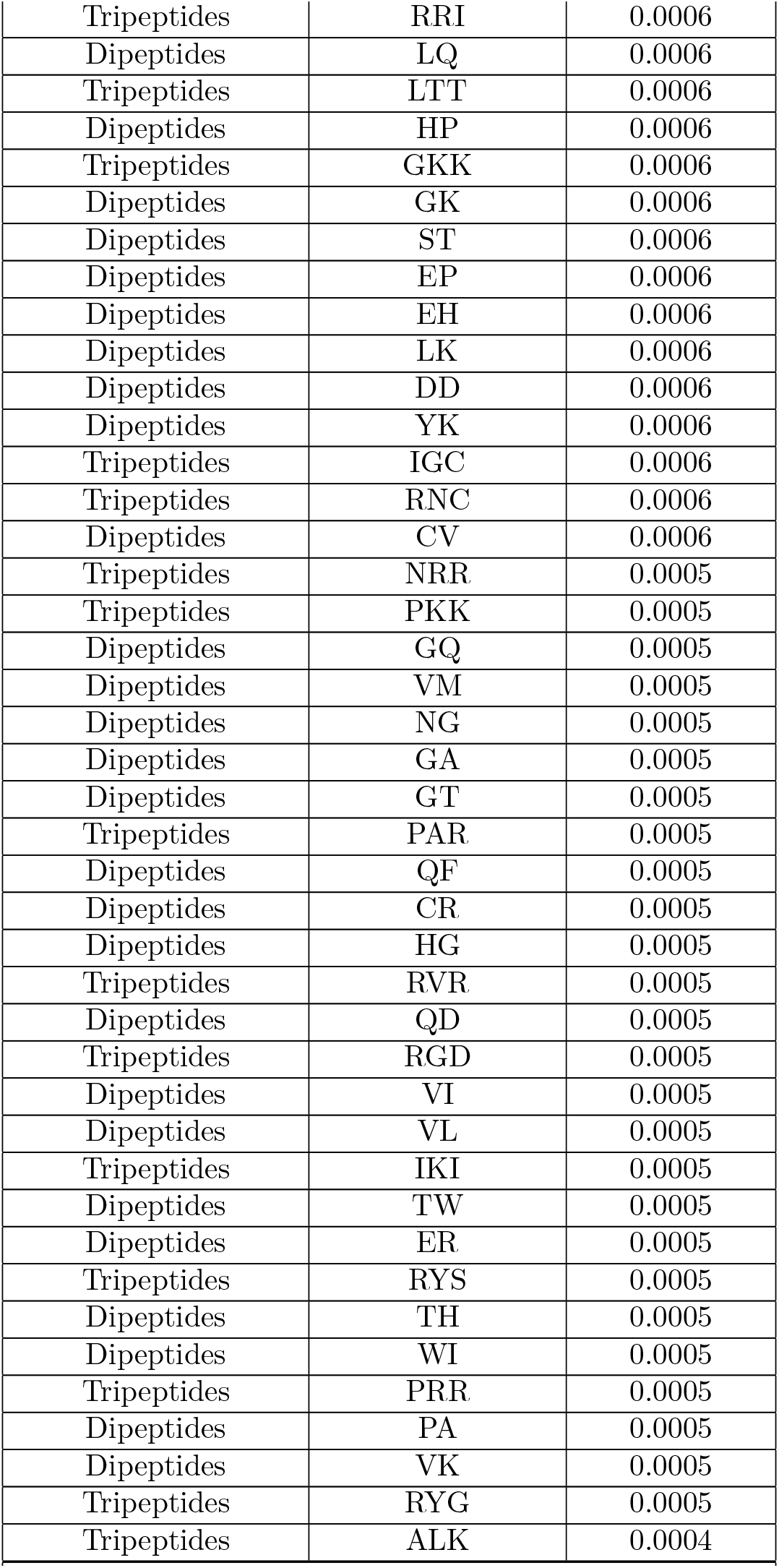
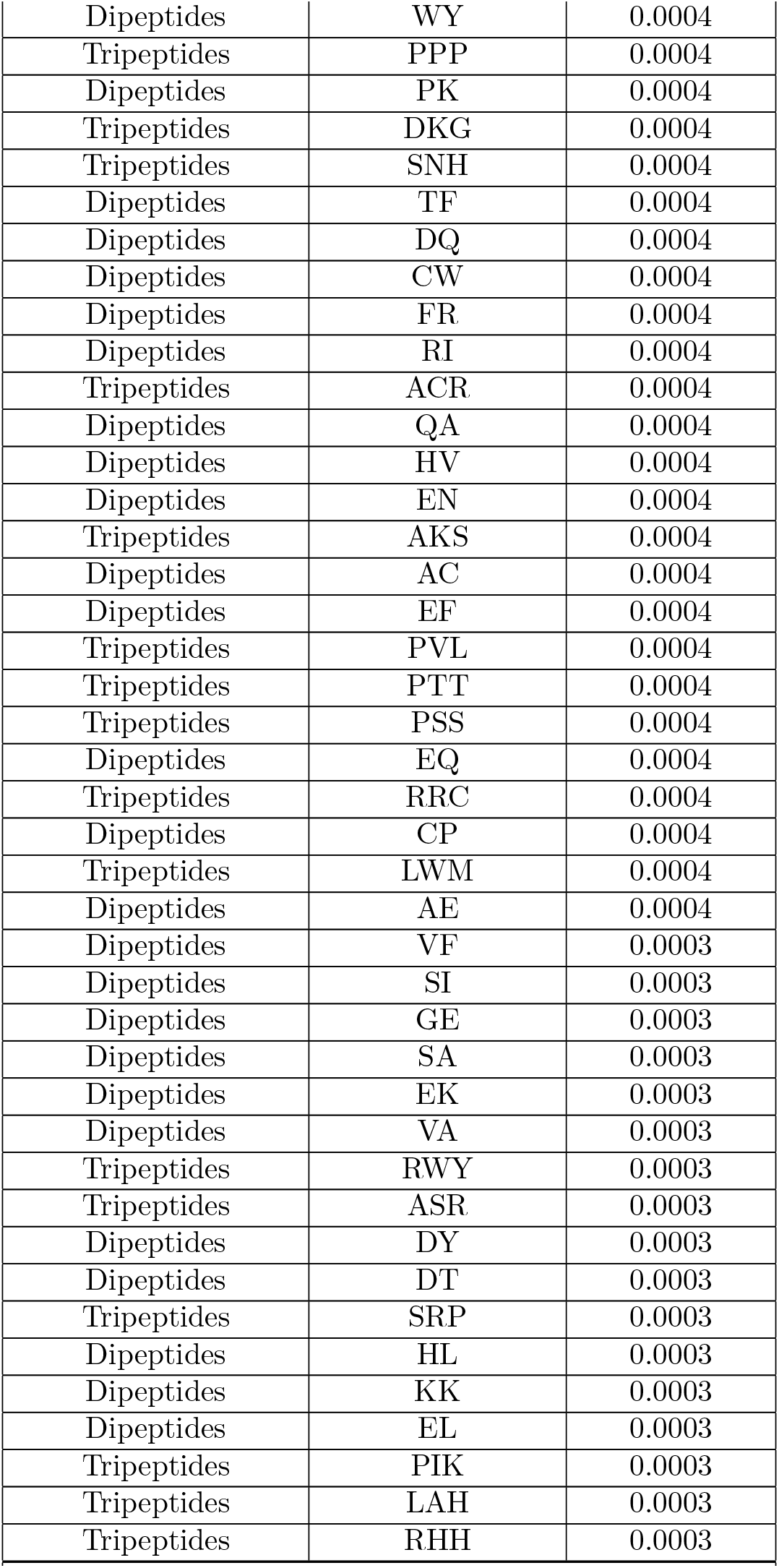
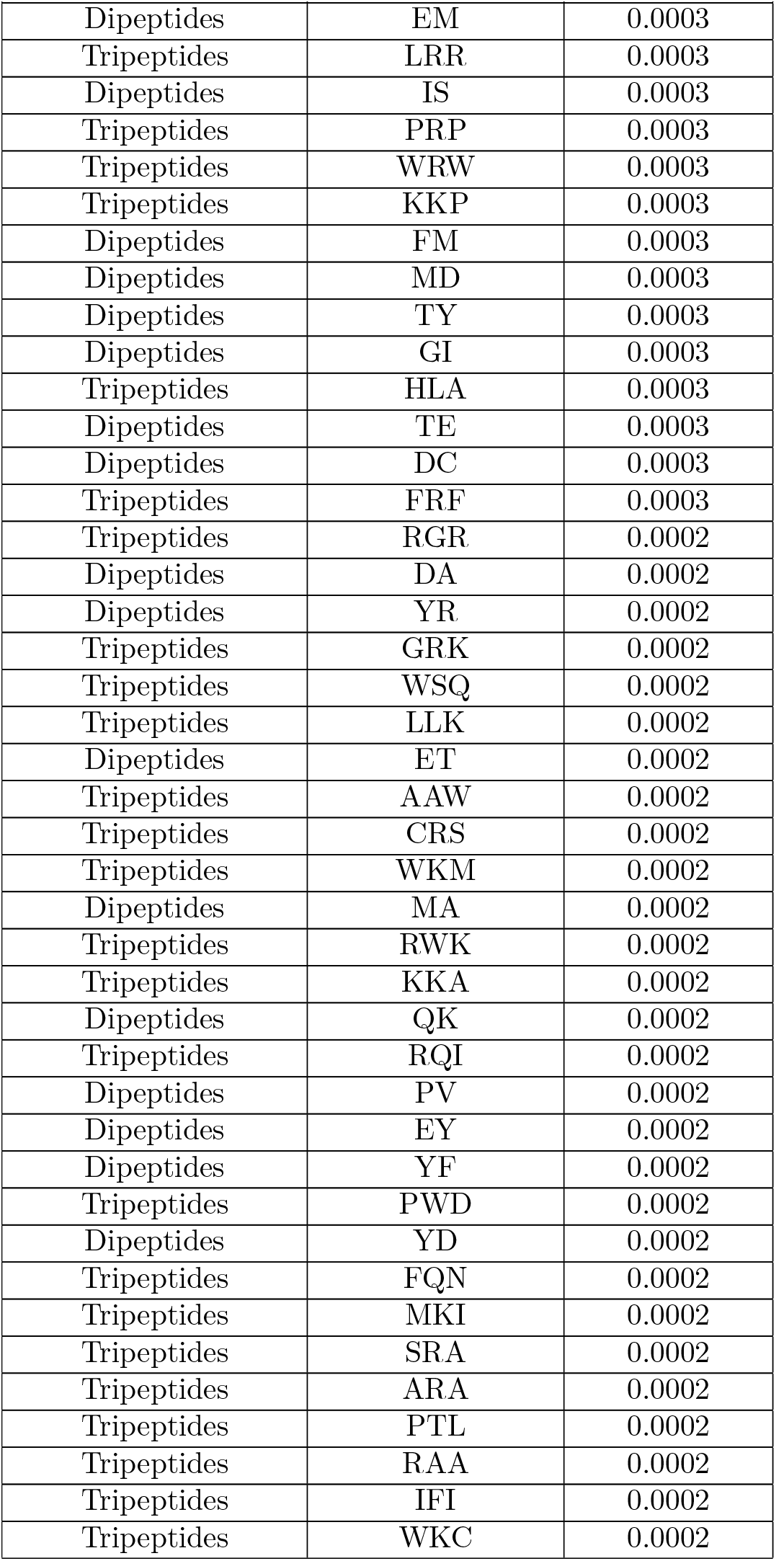
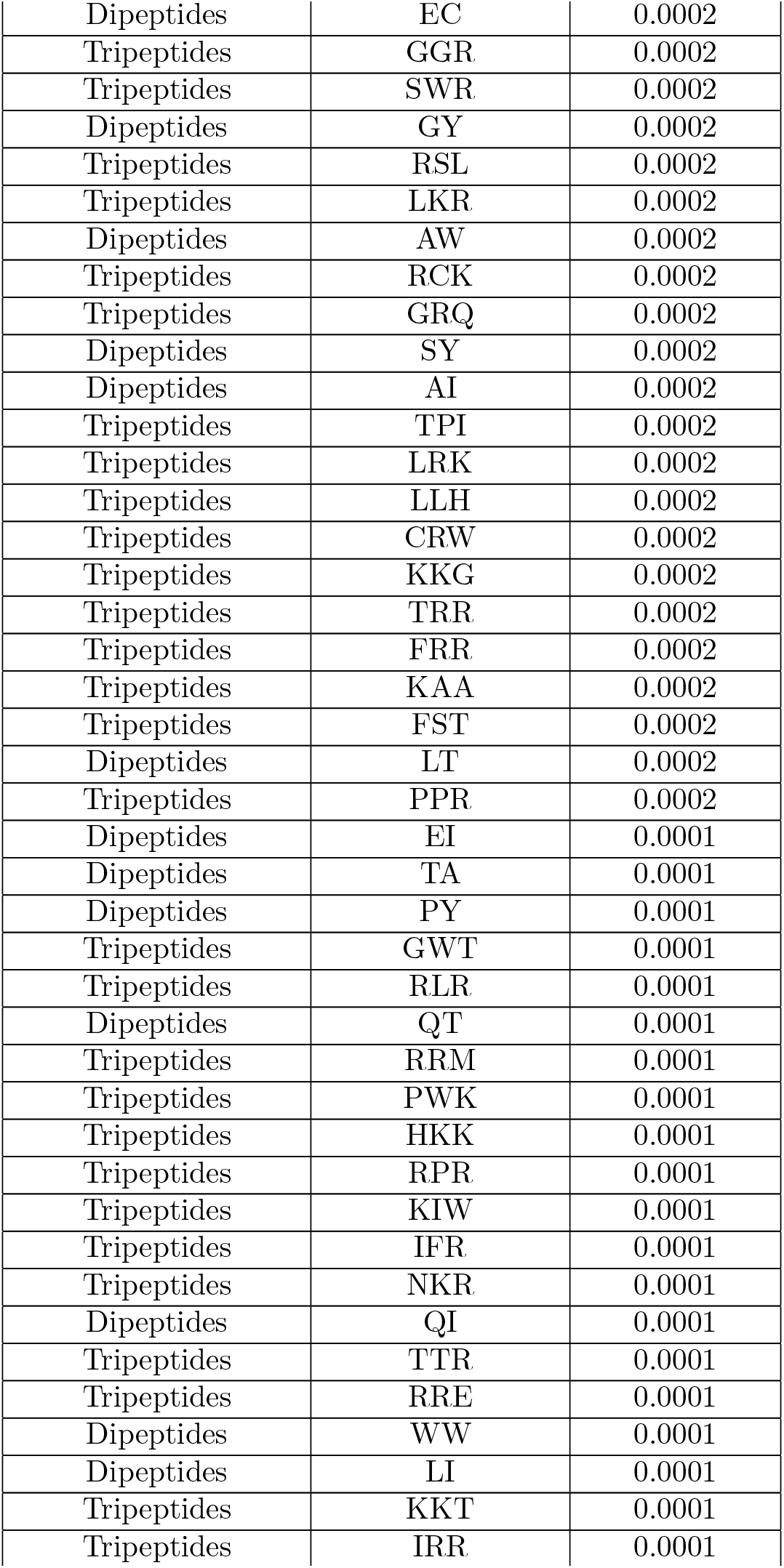
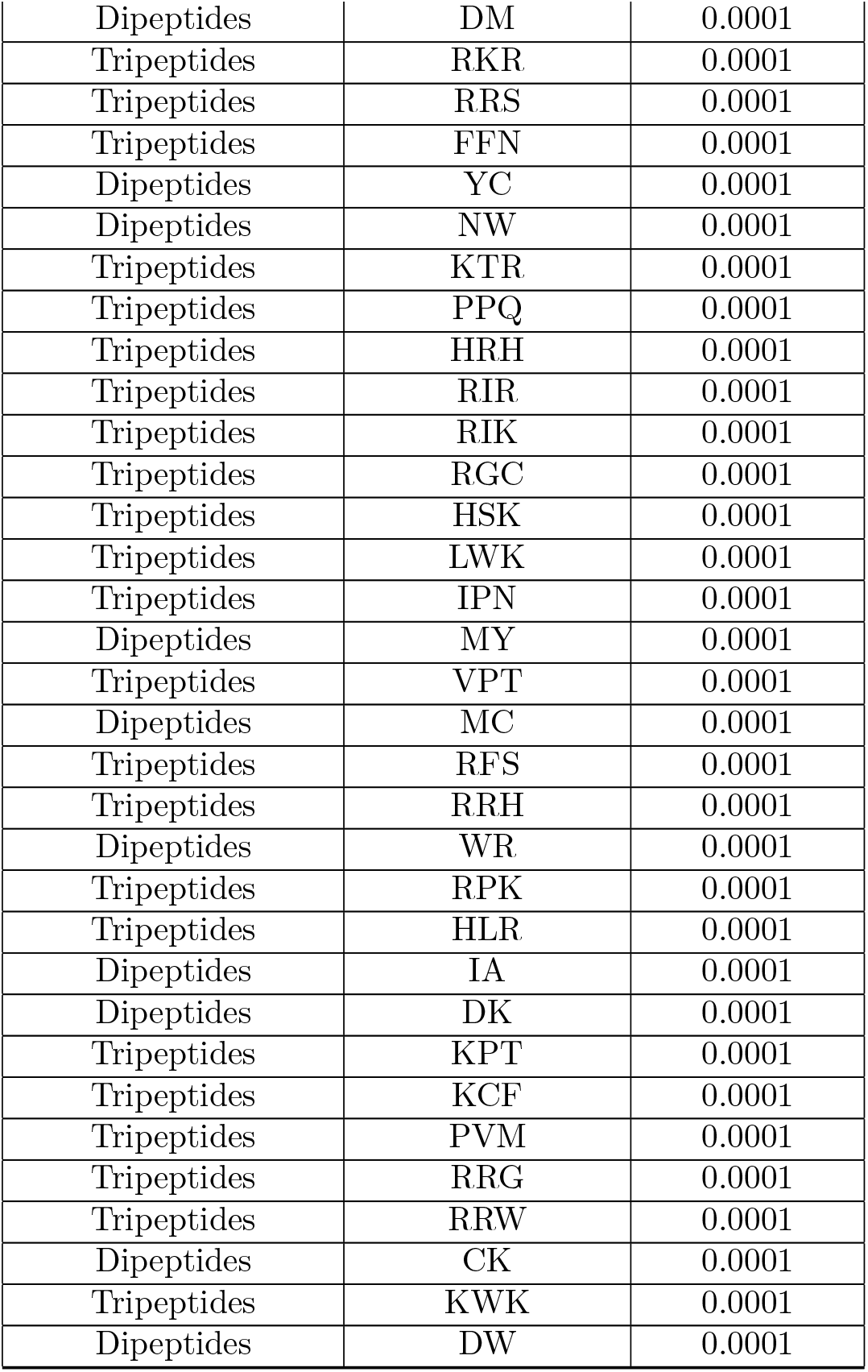
All descriptors.

The t-distributed Stochastic Neighbor Embedding (t-SNE) visualization, a dimensionality reduction strategy used to visualize complex, high-dimensional data, showed a significant separation between CPPs and non-CPPs, as presented in Figure S7. All the TOP20 descriptors were used to plot this t-SNE visualization. Their combination seems to effectively clusters CPPs and non-CPPs, demonstrating their discriminative power. Thus, combining the descriptors discussed above and the small contribuitions of the others 500 features with the ERT algorithm allowed PERSEUcpp to achieve high quality results and out-perform the state-of-the-art competitors.

**Figure S7:**
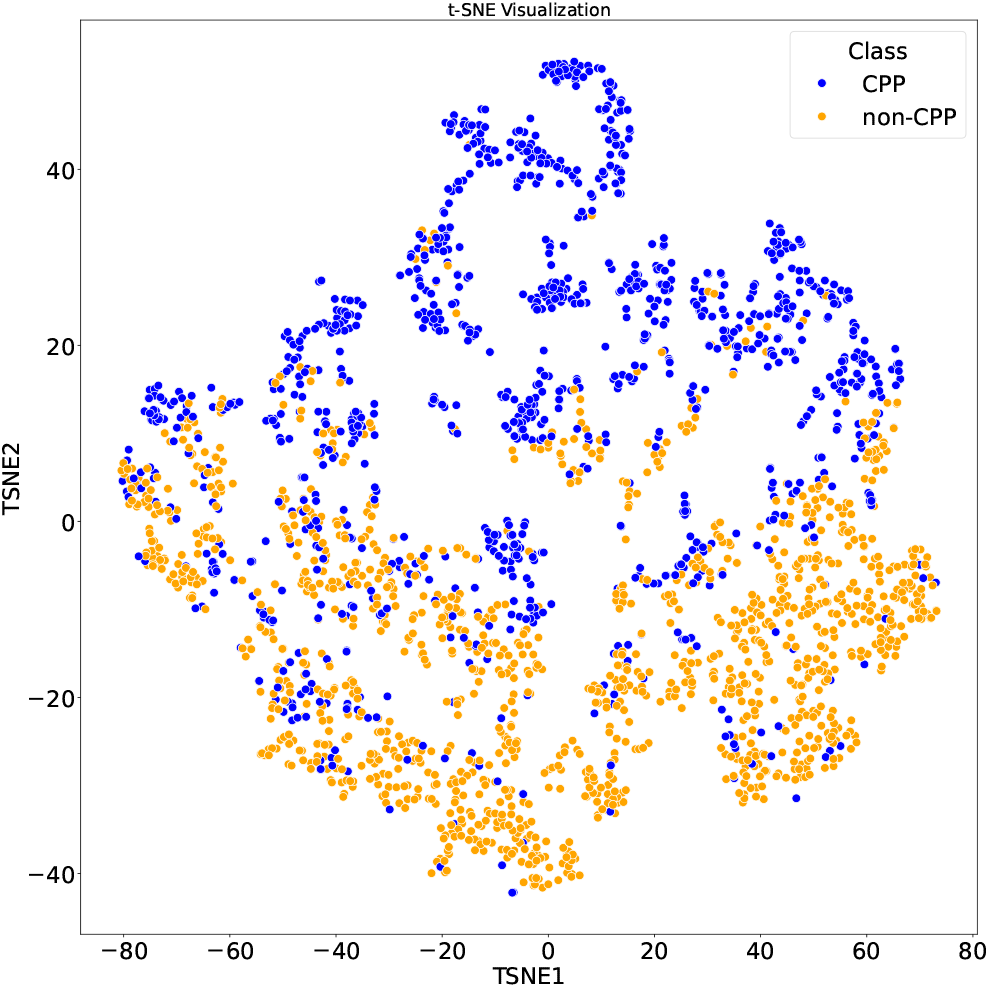
t-SNE visualization of CPPs and non-CPPs. Each point represents a peptide described by a set of calculated descriptors. Points are colored based on their classification: blue for CPPs and orange for non-CPPs. The distinct separation between the blue and orange clusters suggests that the selected features effectively differentiate between CPPs and non-CPPs.

## 4 CPP Efficiency Classifier

### 4.1 Feature importance

**Figure S8:**
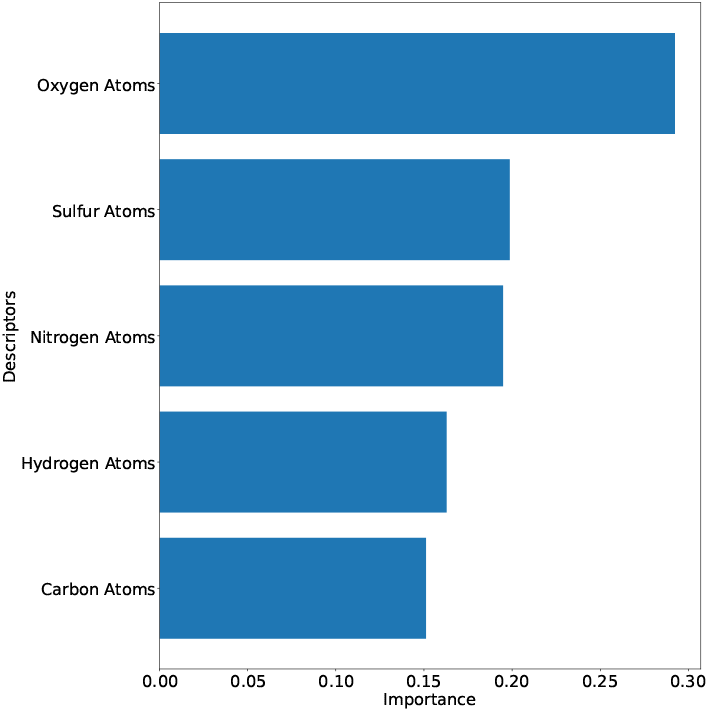
Top features in importance order of each atomic feature: oxygen, sulfur, nitrogen, hydrogen and carbon atoms.

### 4.2 PERSEUcpp compared with state-of-the-art method MLCPP2.0

**Table S13:** Comparison results of the proposed PERSEUcpp. ALL is all normalized features, TRIPEP is the tripeptides group, DIPEP is dipeptides group, CKSAAGP represents the Composition of k-spaced Amino Acid Group Pairs, PQ is the physicochemical group and ATOMIC is the atomic composition group of features.

| Descriptors | MCC | ACC | SN | SP | AUC |
| --- | --- | --- | --- | --- | --- |
| ALL Features | 0.273 | 0.451 | 0.533 | 0.476 | 0.501 |
| TRIPEP | - 0.092 | 0.451 | 0.456 | 0.437 | 0.465 |
| DIPEP | 0.314 | 0.645 | 0.608 | 0.750 | 0.692 |
| CKSAAGP | - 0.052 | 0.580 | 0.695 | 0.250 | 0.565 |
| PC | 0.356 | 0.725 | 0.761 | 0.625 | 0.690 |
| PC + ATM | 0.144 | 0.645 | 0.717 | 0.437 | 0.675 |
| PC + ATM + CKSA | 0.145 | 0.645 | 0.700 | 0.423 | 0.631 |
| ATM + CKSA | -0.032 | 0.564 | 0.652 | 0.312 | 0.559 |
| ATM + DIP | -0.032 | 0.564 | 0.652 | 0.312 | 0.559 |
| ATM + TRIP | -0.096 | 0.419 | 0.391 | 0.500 | 0.559 |

**Table S14:** Comparison results of the proposed PERSEUcpp with TOP-N features of each descriptors group.

| Descriptors | MCC | ACC | SN | SP | AUC |
| --- | --- | --- | --- | --- | --- |
| TOP10 | 0.128 | 0.548 | 0.521 | 0.625 | 0.522 |
| TOP50 | -0.151 | 0.403 | 0.391 | 0.437 | 0.493 |
| TOP100 | -0.054 | 0.483 | 0.500 | 0.437 | 0.497 |
| TOP150 | -0.112 | 0.435 | 0.434 | 0.437 | 0.493 |
| TOP200 | -0.019 | 0.483 | 0.478 | 0.500 | 0.504 |
| TOP250 | 0.019 | 0.516 | 0.521 | 0.500 | 0.513 |
| TOP300 | 0.003 | 0.500 | 0.478 | 0.562 | 0.539 |

